# Nanoformulated Remdesivir with Extremely Low Content of Poly(2-oxazoline) - Based Stabilizer for Aerosol Treatment of COVID-19

**DOI:** 10.1101/2022.01.21.477258

**Authors:** Jacob D. Ramsey, Ian E. Stewart, Emily A. Madden, Chaemin Lim, Duhyeong Hwang, Mark T. Heise, Anthony J. Hickey, Alexander V. Kabanov

## Abstract

The rise of the novel virus SARS-CoV2 which causes the disease known as COVID-19 has led to a global pandemic claiming millions of lives. With no clinically approved treatment for COVID-19, physicians initially struggled to treat the disease and there is still need for improved anti-viral therapies in this area. We conceived early in the pandemic that an inhalable formulation of the drug Remdesivir which directly targets the virus at the initial site of infection could improve therapeutic outcomes in COVID-19. We developed a set of requirements that would be conducive to rapid drug approval: 1) try to use GRAS or GRAS similar reagents 2) minimize excipient concentration and 3) achieve a working concentration of 5 mg/mL Remdesivir to achieve a deliverable dose which is 5-10% of the IV dose. In this work, we discovered that Poly(2-oxazoline) block copolymers can stabilize drug nanocrystal suspensions and provide suitable formulation characteristics for aerosol delivery while maintaining anti-viral efficacy. We believe POx block copolymers can be used as a semi-ubiquitous stabilizer for the rapid development of nanocrystal formulations for new and existing diseases.

## Introduction

The emergence of Severe Acute Respiratory Syndrome Coronavirus 2 (SARS-CoV2) infection initiated a global pandemic of coronavirus disease 2019 (COVID-19)^1^. The infection presents classic symptoms of viral infection such as fever, cough, and fatigue, complicated by acute respiratory distress syndrome (ARDS), which is associated with serious morbidity and mortality^2–4^. With no clinically approved treatment as an initial line of defense, health care workers initially struggled to treat the novel COVID-19. Since the onset of the pandemic, several antiviral agents against SARS-CoV2 have entered clinical trials and/or received authorization for treatment of COVID-19. These drugs target either the TMPRSS2 protease required for viral entry into cells, the RNA-dependent RNA polymerase (RdRp) required for virus replication, or the 3CL protease which is the primary coronavirus (CoV) protease required for processing the CoV nonstructural polyprotein^5,6^. The first approved drug was Gilead’s Remdesivir (GS-5734) which targets the RdRp and appeared the most promising candidate to treat COVID-19. On May 1, 2020 the US Food and Drug Administration (FDA) authorized the emergency use of Remdesivir and it gained full FDA approval in October 2020. Until recently, Remdesivir which is administered as intraveneous (IV) infusion was the only small molecule intervention in which evidence supported benefits for both time to symptom resolution and duration of mechanical ventilation^7,8^. Although benefits of Remdesivir on mortality has long remained uncertain^9–11^, the drug has shown strong efficacy in high risk outpatients with COVID-19 and reduced risk of hospitalizations by 87% compared to placebo^12^. Most recently, on December 23, 2021 another drug, Molnupiravir (MK-4482) from Merck, which also targets RdRp, has recieved an Emergency Use Authorization (EUA) for situations where other FDA-authorized treatments for COVID-19 are inaccessible or are not clinically appropriate (https://www.fda.gov/news-events/press-announcements/coronavirus-covid-19-update-fda-authorizes-additional-oral-antiviral-treatment-covid-19-certain). The clinincal trial of Molnupiravir in high risk populations has demonstrated that the drug decreases the risk of hospitalization and severe disease by up to 50% compared to placebo^13–15^. Even more impressive results were seen with Pfizer’s Paxlovid (3CL protease inhibitor nirmatrelvir (PF-07321332) and ritonavir tablets, co-packaged for oral use) which was granted EUA on December 22, 2021 for the treatment of mild-to-moderate COVID-19 in adults and pediatric patients (https://www.fda.gov/news-events/press-announcements/coronavirus-covid-19-update-fda-authorizes-first-oral-antiviral-treatment-covid-19). Paxolovid significantly reduced the proportion of people with COVID-19 related hospitalization or death from any cause by 88% compared to placebo among patients treated within five days of symptom onset and who did not receive COVID-19 therapeutic monoclonal antibody treatment. Both Molnupiravir and Paxlovid are given orally and can be taken outside a hospital setting. This would allow for the early treatment of COVID-19, which is a critical breakthrough in the treatment of COVID-19^13,16^. Additionally, there is an antibody cocktail from Regeneron which has obtained EUA, however this can only be delivered in a hospital setting^17–19^. As the unprecedented effort to contain and control COVID-19 has had varying success, new variants continue to emerge. The recent surge of the COVID-19 cases in the United States and worldwide due to the spread the Omicron variant of SARS-CoV2 is most concerning. The need for relief remains urgent and a priority for the World Health Organisation (WHO) as immunization efforts in many counries have been much slower than desired.

There is room for improvement of the current Remdesivir formulation, with some disputing its efficacy^10,11,20^. Gilead’s current formulation of Remdesivir is administered parentally with the aid of sulfobutyl ether beta cyclodextrin ((SBECD); 5 mg/mL Remdesivir at 1:30 Remdesivir:SBECD weight ratio)^8^. Gilead has shown no significant difference between a 5 day and 10 day course of Remdesivir therapy^21^. There are many challenges facing the delivery of Remdesivir to the lungs, the primary disease site. Ideally, an oral dosage form would be ideal and allow for administration of Remdesivir outside of a clinical setting reducing hospitalizations and exposure of health care workers. However, this is not possible due to Remdesivir’s intrinsic liver toxicity^20^. The first pass effects after absorption from the GI tract would mean very high concentrations of Remdesivir in the liver causing undesirable toxicity. Additionally, the use of an excipient based solubilizer means that a large portion of this hydrophobic drug will partition to the protein bound fraction in circulation limiting the unbound fraction which is capable of pharmacodynamic activity in the lungs. The clearance rate from the body will be fast and the drug can distribute itself to many off target sites^20^. Some have argued that the use of the parent nucleoside, GS-441524, would be better suited for treatment as Remdesivir is rapidly converted to the parent nucleoside *in vivo* anyways^20^. However, if delivered directly to the target site, Remdesivir might be a better choice than the parent nucleoside. Lung delivery presents no concern for traditional first-pass metabolism, and there are few drug metabolizing enzymes to convert Remdesivir back to the parent nucleoside prior to cell uptake. Esterases are ubiquitously expressed in the lungs, although with some degree of variability, and can readily convert Remdesivir to the monophosphate form and then subsequent phosphorylation can activate the drug for RdRp inhibition^22,23^. Remdesivir requires multiple metabolic steps to reach its bioactive tri-phosphate form. Recently, Gilead has utilized their same Remdesivir formulation as an aerosol for delivery directly to the lungs^24^. Utilizing a delivered dose which is roughly 5% of the IV dose, they demonstrated comparable efficacy to the IV formulation in a non-human primate SARS-CoV2 model. To minimize excipient levels in our formulation, we focused on developing a polymer-stabilized nanoparticle formulation for Remdesivir for aerosol delivery directly to the lungs.

## Materials and Methods

### Materials

Methyl Triflate, 2-Methyl-2-oxazoline, 2-*n*-Butyl-2-oxazoline, Acetonitrile, Ether, Acetone, Deuterated Water, 3.5 kDa Dialysis membrane, Potassium carbonate, Methanol, Tyloxapol, Oleic Acid, Piperidine, Ultra Pure Water, phosphate-buffered saline (PBS), and 3.5 kDa Slidealyzers were purchased from Sigma Aldrich. Remdesivir and Tyloxapol were purchased from Medkoo, Pluronic F127 (F127) was obtained from BASF. Sulfobutylether-β-cyclodextrin (SBECD) was purchased from Medchem Express. A Salter Labs 8900 jet nebulizer (SunMed; Grand Rapids, MI) was used for all aerosol studies. Impaction stages were washed with methanol (Fisher Scientific).

### Polymer Synthesis

Triblock copolymer P[MeOx-*b*-BuOx-*b*-MeOx], P2 with degrees of polymerization of the blocks 38-33-38 was synthesized according to previously described methods^25,26^. For the synthesis of polymers, methyl tri-fluoromethanesulfonate (MeOTf), 2-methyl-2-oxazoline (MeOx), and 2-*n*-butyl-2-oxazoline (BuOx) were dried by refluxing over calcium hydride (CaH_2_) under inert nitrogen gas and subsequently distilled. Briefly, MeOTf was added to a reaction flask under low oxygen and water vapor conditions and in argon gas environment. 3 mL of dry acetonitrile was used as a solvent. MeOx was added at desired molar ratio and mixed overnight at 80 °C. Complete monomer consumption and block length were confirmed by proton nuclear magnetic resonance (^1^H-NMR) (methanol solvent). BuOx was added to the reaction mixture under dry conditions at the desired block length and mixed overnight at 80 °C. ^1^H-NMR was again used to confirm completion and block length. MeOx was then added and the reaction mixture was stirred overnight at 80 °C. Terminating Piperidine was added in 3-fold molar excess and the reaction mixture was mixed overnight at room temperature. Potassium Carbonate was added to dry the mixture. The mixture was then gravity filtered and washed with acetone. Acetone was removed by Rotovap and then the mixture was added to ice cold ether in a 9:1 ether:reaction mixture volumetric ratio. The vials were then centrifuged at 1000xG for 5 minutes to pellet the precipitated polymer. Ether was decanted and the precipitate was dissolved in ultrapure water and dialyzed (3.5 kDa membrane) against water for 4 days changing the surrounding medium every 24 hours. Samples were lyophilized and final structure confirmed by ^1^H-NMR.

### Bottom-Up Nanocrystal Synthesis

Remdesivir nanocrystals were prepared by the solvent-anti solvent precipitation method^27^. Briefly, Remdesivir was solubilized in methanol at a concentration of 80 mg/mL and polymers (F127, P2, Tyloxapol) were dissolved to a concentration of 10 mg/mL in methanol. Oleic acid was used at a working concentration of 1 mg/mL in methanol. Nanocrystal suspensions were synthesized at Remdesivir concentrations ranging from 2.5-20 mg/mL. The desired volume of suspension in UltraPure Water (Ranging from 2-10 mL in this work) was placed in a scintillation vial under rapid stirring (900 rpm). Remdesivir and polymer stabilizers were mixed together in the desired ratios with a given weight % of stabilizer as determined by **Equation 1**. For example, in a 2 mL, 5 mg/mL suspension, 125 μL of Remdesivir and 50 μL of P2 would be used. These were added to the scintillation vial under rapid stirring and allowed to mix for 30 seconds. The suspensions were then transferred to microcentrifuge tubes and centrifuged at 10000 x G for 5 minutes to pellet most of the suspension. Supernatant was removed and the nanocrystals could then be redispersed to the desired concentration. For long term storage, samples were frozen in liquid nitrogen with a concentration of 20% trehalose (v/v) as a cryopreservant.

**Equation 1:** Mass Stabilizer/Mass Remdesivir*100

### SBECD Formulation Preparation

Remdesivir Inclusion complexes (IC) were prepared by adding Remdesivir in methanol dropwise at roughly 7 drops/minute to an aqueous solution of SBECD under vigorous stirring at a 1:30 Remdesivir:SBECD ratio by mass creating a roughly 2 mg/mL Remdesivir solution. After drops finished, the solution was stirred for an additional 1 hour. The solution was then put on Rotovap to remove methanol and then lyophilized to form a powder which is 3.2% Remdesivir by weight. Solutions were hydrated to the desired concentration in ultra-pure water and allowed to stabilize for 15 minutes before use.

### Nanocrystal Size and Zeta Potential Characterization

Nanocrystal particle sizes were measured by Dynamic Light Scattering (DLS) where we used both the intensity distribution and the number-based distribution measurements to gain insights about the formulation. Morphology was determined using Transmission Electron Microscopy (TEM). Zeta Potential of nanocrystals were measured using a Particle Metrix Zetaview Instrument at 20-fold dilution in 0.1X PBS.

### In Vitro Drug Release Studies

Remdesivir nanocrystals and SBECD IC’s were prepared at 2 mg/mL and 5 mg/mL and the release rate from the formulations was measured under sink conditions in PBS at 37°C. 100 uL of sample was placed in a 3.5 kDa slidealyzer dialysis devices after washing the membrane with PBS. Samples were incubated in triplicate at time points of 0.5, 1, 2, 4, 8, and 24 hours. The solution was removed from the dialysis device and lyophilized. It was then rehydrated in 100 μL of methanol and drug concentration was measured by HPLC after centrifugation and removal of salts.

### HPLC Analysis of Remdesivir

Remdesivir concentrations were analyzed by Agilent 1100 HPLC system on a Reverse Phase C18 column from Supelco. Briefly, 70 μL of Remdesivir to be measured in methanol was added to 30 μL of HPLC water for sample analysis. Mobile Phase was 70:30 methanol:water ratio with a 1.0 mL/minute flow rate. Column temperature was set to 40 °C and the retention time of Remdesivir was about 5 minutes. Injection volume was 10 μL. Remdesivir was analyzed at 280 nm on a UV Photodiode Array detector.

### Methanol Detection by NMR

For quantifying methanol removal from the nanocrystal suspensions, ^1^H-NMR was used. ^1^H-NMR spectra were recorded on an INOVA 400 at room temperature. The spectra were calibrated using the solvent signals (D_2_O 4.80 ppm). Samples of nanocrystal suspensions were dissolved in deuterated water before and after centrifugations. Samples were centrifuged to pellet Remdesivir nanocrystals and then supernatant was analyzed on ^1^H-NMR. The methanol solvent peak area near 3.3 ppm was used to determine the amount of methanol remaining in the nanocrystal suspension.

### In Vitro Antiviral Studies

Antiviral studies were performed according to previous protocols^28,29^. The A549 cells expressing Ace2 (A549-Ace2) were seeded at 1 × 10^4^ cells/well in black walled clear bottom 96 well plates the day prior to treatment and infection. Drug stocks and controls were prepared fresh within an hour of each replicate and diluted in saline prior to addition to the 96 well plates. Prepared drug stocks were then diluted 1:100 in cell maintenance media (Gibco DMEM supplemented with 10% heat-inactivated FBS, 1% Gibco NEAA, 1% Gibco Pen-Strep) to achieve a 2x concentration. Maintenance media was removed from cells and cells were pre-treated for 1hr with 2x drug. Cells were then either mock infected or infected at a Multiplicity of Infection (MOI) = 0.02 with SARS-CoV-2-nLuc. The final concentration of drug after the addition of inoculum was 1x. Cells were then incubated for 48 hr at 37°C with 5% CO_2_. Virus growth was measured with NanoGlo Luciferase Assay System (Promega) and cell viability was measured with CellTiter-Glo 2.0 Cell Viability Assay (Promega). Luminescence was measured using a Promega GloMax Explorer System plate reader. Background luminescence from vehicle treated wells subtracted from all other treated wells. Drug treatment experiments were performed in three independent experiments in technical duplicate.

### Aerosol Generation Inertial Impaction and Aerodynamic Characterization

Inertial impaction was performed using a Next Generation Impactor (NGI; MSP Corp.; MN, USA) following United States Pharmacopeia (USP) method <1601>. Impactor stages were precooled to 4 °C for at least 90 minutes before experiments. Vacuum for the NGI was set to 15 L/min and a solenoid wired to the vacuum pump was set to 2 minutes. Aliquots of the Remdesivir formulation were synthesized and frozen at 10 mg/mL, thawed and then diluted to 1.5 mg/mL. The nebulizer (Nebutech 8960) was filled with 3 mL of the Remdesivir formulation and affixed to the inlet of the NGI via custom mouthpiece adapter. The nebulizer was actuated by connecting the pressurized air tubing (50 psi)and generating the aerosol at an airflow rate of 8L/min. Next the solenoid was switched on and nebulized material was collected over the 2 minutes. This was repeated thrice with fresh NGI plates, inlets, and Remdesivir nanocrystal suspensions.

The impactor size segregates the aerosol with cut-off aerodynamic diameters of A, B, C, D, E, F, G μm for stages 1-7, respectively, from which the particle size distribution can be constructed (USP<1601>). USP Impactor Cutoffs at the specified conditions are as follows: Stage 1 = >14.1 um, Stage 2 = <14.1 um, Stage 3 = <8.61, Stage 4 = <5.39 um, Stage 5 = <3.3 um, Stage 6 = <2.08 um, Stage 7 = <1.36 um, and Stage 8 = <0.98 um. The deposited aerosol on the inlet, stages 1-7, and multi-orifice filter (MOC) of the NGI was assayed by HPLC as described above to quantify Remdesivir mass. The values were used to construct an aerodynamic particle size distribution (APSD) from which mass median aerodynamic diameter (MMAD), geometric standard deviation (GSD) (**Equation 2**), and fine particle dose (FP Dose; mass particles < 5.39 μm at 15 L/min) could be determined. MMAD was calculated by plotting the cumulative mass percent undersize of the deposited Remdesivir against its corresponding stage effective cutoff diameter, translating to a probability (PROBIT) vs. logarithm effective cutoff diameter plot, and fitting a line through points on either side of 50^th^ percentile mass collected and solving for particle size (equal to 0 on PROBIT scale). GSD was similarly calculated by determining the particle size at the 84^th^ and 16^th^ percentile (1 and −1, respectively, on PROBIT scale) and taking the square root of the former divided by the latter^30^. The FP Dose was determined by summing the mass Remdesivir collected on stages 4 through MOC of the NGI. The FP Dose % was determined by the percentage of delivered Remdesivir which was in the respirable fractions. The percent of the suspension output is the mass of Remdesivir delivered divided by the nominal dose.

**Equation 2:** GSD = [% mass of distribution at D_84_/% mass of distribution at D_16_]^0.5^

### ^1^H-NMR of Polymers

^1^H-NMR spectra were recorded on an INOVA 400 at room temperature. The spectra were calibrated using the solvent signals (MeOD 3.31 □ppm).

## Results

In late March 2020, we conceived that the delivery of an aerosol formulation of Remdesivir directly to the lungs could improve therapeutic outcomes in COVID-19^31,32^. By directly targeting the site of disease, the lungs, one could improve therapeutic outcomes while lowering both the overall dose and amount of excipient to potentially decrease negative side effects, like avoiding liver toxicity and systemic activity at sites unrelated to the disease pathology. To achieve this, we generated a set of requirements which could be conducive to the rapid approval of an aerosol formulation of Remdesivir. First, we estimate the targeted concentration of 5 mg/ml for Remdesivir in the aqueous dispersion for aerosol preparation to have a feasible volume for nebulizer administration. We assessed that we would only need to deliver between 10-20% of the current Remdesivir dose, which is an IV infusion (200 mg loading, 100 mg maintenance). Thus, as an aerosol, we would only need to deliver 10-20 mg. Second, we proposed that the amount of the excipient in this dispersion should not exceed 5% of the total drug weight (≤ 0.25 mg/ml excipient). This “stringent” approach to the formulation design with the excipient being its minor component, is advantageous for regulatory approval as it minimizes safety concerns. Finally, we propose using generally regarded as safe (GRAS), “GRAS Similar” (such as pluronics, which contain the GRAS PEG block), or excipients used in previously approved drug products. For nebulized aerosols for inhalation, it is also important to consider factors such as solution foaming and aerosolized droplet size^30,33,34^. The injectable form of Remdesivir is not ideal for inhalation due to the presence of significany solubilizing excipient (5 mg/ml drug and 150 mg/ml SBECD solubilizer)^8^. We postulate that the nanoparticle approach will allow preparation of aqueous dispersion of Remdesivir with the desired drug concentration, low content of the excipient, and suitable rheological and surface properties.

There are two major approaches to nanocrystal preparation: top-down and bottom-up approaches. Top-down approaches involve wet-milling processes, which is a commercially scalable process offered by several providers^27,35^. We opted to pursue the laboratory scale bottom-up, solution/precipitation-based approach to try and control the growth of Remdesivir drug crystals into nano-sized particles and determine the optimal composition of the nanocrystal suspension that address our chosen parameters. In this study, we explored factors such as mixing type (bulk mixing by stir bar vs. homogenization), solution temperature (4 °C vs. room temperature), drug concentration, organic solvent content, stabilizer composition, and the order of reagent addition. Once optimized, we explored the stability of the formulation with respect to duration of longterm storage in addition to multiple metrics of *in vitro* evaluation.

Initially, we established a list of common excipients used to stabilize nanocrystal formulations some of which are excipients used in product approved for clinical use and others were GRAS or “GRAS similar” compounds^36^. These reagents are shown in **Supplemental Table S1** with some associated characteristics of the excipients.

Of note, the hydrophilic-lipophilic balance (HLB) value of some surfactant excipients used herein as stabilizers is quite high, which is known to cause foaming in solutions. These stabilizers were explored in various facets, beginning with a poloxamer F127, because it is a GRAS similar compound. F127 is an amphiphilic A-B-A triblock copolymer consisting of flanking hydrophilic blocks of poly(ethylene oxide) (PEO) and a central hydrophobic poly(propylene oxide) (PPO) block. PEO, also known as poly(ethylene glycol) (PEG). PEG is a GRAS agent commonly used as a sterically stabilizing molecule in nanoparticle formulations^37,38^. Poloxamers have been used in multiple clinically approved products or those evaluated in clinical trials including gel, micelle, and topical formulations. F127 in particular, was in a Phase II polymeric micelle injectable formulation^39^.

To prepare F127 stabilized Remdesivir nanocrystals, F127 was dissolved in water at concentrations ranging from 0.0 to 0.25 mg/ml (which corresponds to 0-5% by weight of Remdesivir e.g. 5% F127 is 0.25 mg/mL F127 compared to 5 mg/mL Remdesivir). Remdesivir in methanol (80 mg/mL) was then added to the solution to make it 5.0 mg/mL in 2 mL water. This synthetic procedure is depicted graphically in **Supplemental Figure S1**. The solution was stirred rapidly, and the growth of the nanocrystals was monitored over time by DLS. **Figure 1A** shows the effect of the starting stabilizer concentration on the size of the Remdesivir nanocrystals. As we expected, the lower amounts of F127 yielded higher nanocrystal sizes as there were more Remdesivir molecules per molecule of F127. In the absence of F127, large multi-micron sized particles were formed immediately that irreversibly precipitated out of solution soon after. In the presence of 0.25 mg/ml F127 (5% of Remdesivir by weight), the nanocrystals formed with effective diameter of 341.1 nm, as measured by DLS (**Figure 1A**). The stability of these nanocrystals was monitored over a period of 5 days where their size distribution shifted towards larger particles-perhaps due to an ostwald ripening process. Eventually, they aggregated completely and precipitated out of solution irreversibly after the 5^th^ day (could not be measured on DLS) (**Figure 1D**). However, there was still methanol in this solution, which precludes the use of this composition as a therapeutic. To eliminate methanol from the solution, and perhaps improve stability, we lyophilized the compositions both with and without 9% sucrose and then rehydrated the powders with ultrapure water to the same original volume of the lyophilized aliquot. Sucrose is a known cryoprotectant which could confer stability to the lyophilized formulation^40–43^. Although the particles in the sample lyophilized with sucrose were smaller than those formed in the absence of sucrose (**Figure 1B**) they still were stable in dispersion only for about 30 min after hydration. Some samples of these F127 nanocrystals were prepared and tested as an aerosol immediately post hydration (**Supplemental Figure S2**). Solution foaming led to inconsistent drug output rates over 12 mins (**Supplemental Figure S2A**). Despite these inconsistent output rates, the MMAD remained below or near 5 μm (**Supplemental Figure S2B**), which is at the upper end of what is generally considered respirable (typically defined as particles with MMAD 1-5 μm^44–46^). To increase the consistency of drug output, we tested additional combinations of stabilizers by adding Tyloxapol, oleic acid, and Pluronic L61 in select combinations which could provide anti-foaming effects **(Supplemental Table S2)**. They also did not offer additional nanocrystal stability to enhance the long-term storage needed for clinical translation-solutions still precipitated soon after being hydrated post-lyophilization.

**Figure 1:**
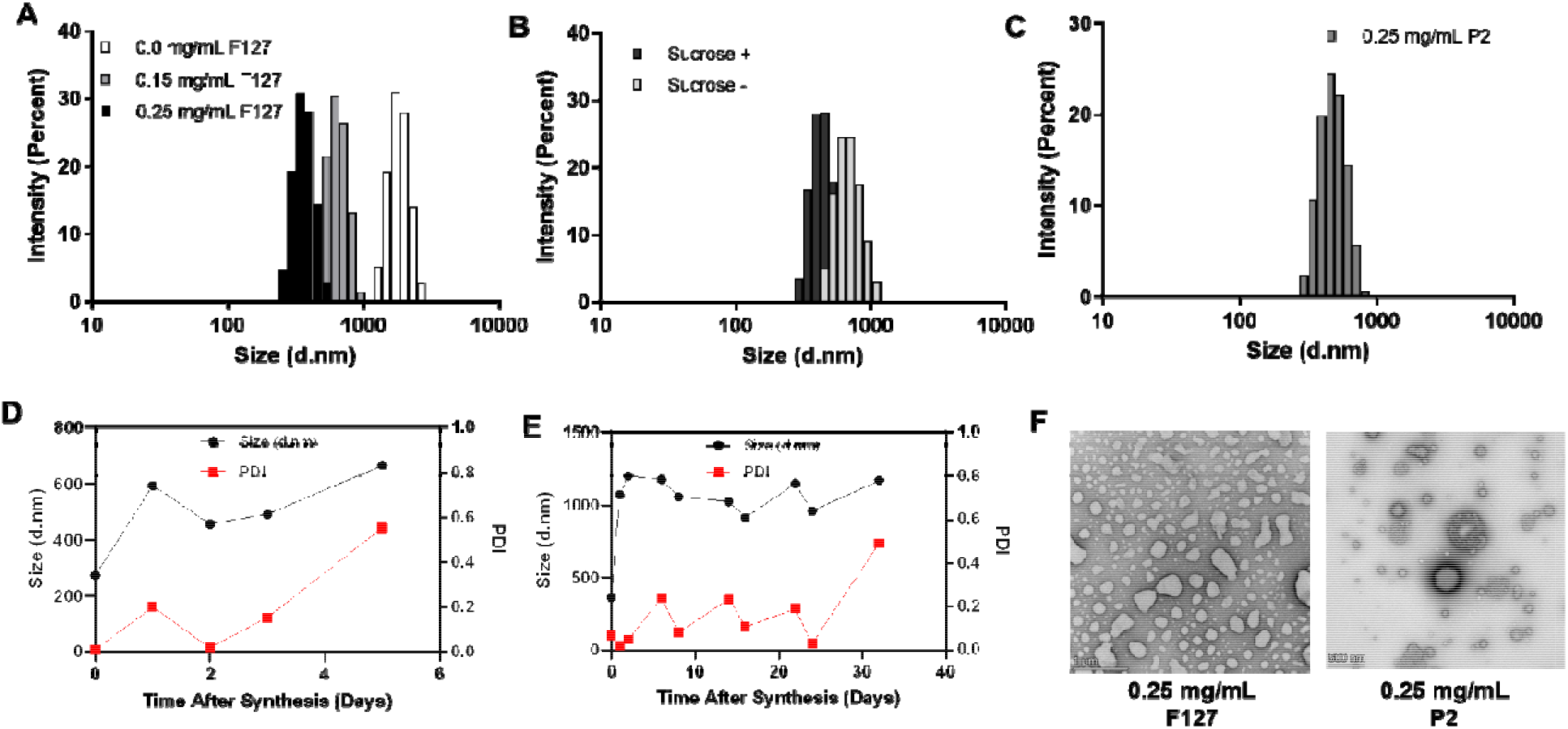
Initial preparation of 5 mg/mL Remdesivir nanocrystals with F127 and P2: **(A-C)** DLS intensity size distribution of nanocrystals after 10 minutes stabilized with **(A)** with various amounts F127 (0.0, 0.15, and 0.25 mg/mL), **(B)** 0.25 mg/mL F127 nanocrystals redispersed after lyophilization with and without 9% sucrose (w/v), and **(C)** 0.25 mg/mL P2. **(D,E)** DLS size by intensity distribution and PDI of nanocrystals stabilized with **(D)** 0.25 mg/mL F127 assessed over 5 days and **(E)** 0.25 mg/mL P2 assessed over 30 days. **(F)** TEM images of nanocrystals.

Ultimately, we determined that the 30-minute stability of the F127 nanocrystals after lyophilization was unsatisfactory. Gilead’s current formulation states that it needs to be used within 24 hours of hydration when stored at room temperature or within 48 hours when stored at 4°C (https://www.gilead.com/-/media/files/pdfs/remdesivir/eua-fact-sheet-for-hcps.pdf?la=en&hash=D4229149DCD2FF6B7E83F4062C4601BB, *Accessed January 7, 2022*)). For our inhalable formulation, we selected 4 hours as a target for the reasonable stability of a formulation. In **Figure 1C** we tested the ability of a novel nanocrystal stabilizer, called P2, which is an A-B-A triblock copolymer comprising a POx backbone. **Supplemental Table S1** shows the structure of P2 block copolymer and **Supplemental Figure S3** has the ^1^H-NMR Confirmation of structure. It has two flanking hydrophilic MeOx A blocks and a central, hydrophobic BuOx B block: P[MeOx-*b*-BuOx-*b*-MeOx]. Notably, the MeOx block is more hydrophilic than PEO in F127, and the BuOx block contains both hydrophobic as well as polar moieties that are capable of ion-dipole interactions and forming hydrogen bonds with drug molecules^47–49^. We posited that the P2 could adsorb at the surface of the growing Remdesivir nanocrystals with the aid of the BuOx block, while the MeOx block could form a protective hydrophilic brush stabilizing the nanocrystals in dispersion. The nanocrystals formed in the presence of 0.25 mg/ml P2 were larger than those synthesized with F127, near 400-500 nm, but were stable in solution for greater than 25 days after synthesis (**Figure 1E**). TEM of the F127 and P2 stabilized nanocrystals is seen in **Figure 1F**. Additional tests using the excipient Tyloxapol, which is included in a clinically approved aerosol product, did not improve nanocrystal size or stability over those of F127 **(Supplemental Table S2)**. Because of this, we ultimately elected to proceed with the P2 stabilized nanocrystals as our flagship formulation for further analysis. Initial observations during nebulization indicated little to no foaming which could impact output over time.

After determining that the P2 stabilized nanocrystals could be prepared at suitable sizes for aerosol delivery, with minimal excipient, and nebulized with minimal foaming, we proceeded to evaluate the stability of the solution during the process of methanol removal. We made one small change to the synthetic procedure depicted in **Supplemental Figure S1**. In the new procedure, we mix both Remdesivir and P2 in methanol and add the solution to ultrapure water under mixing. At this point, we wanted to evaluate the feasibility of multiple storage conditions for the P2 stabilized nanocrystals, namely, freezing and lyophilization. For the freezing process, we removed methanol by a series of centrifugations. We quantified the remaining methanol using ^1^HNMR spectroscopy. We monitored the size and PDI of the nanocrystals after two centrifugations and noted in **Supplemental Table S3** that the size was maintained mostly in the 400-450 nm range prior to freezing and a low PDI of about 0.20 and below was maintained as well. Nanocrystals were readily resuspended into solution after pelleting indicating their stability through this process. From this, we concluded that the nanocrystals were indeed stable through centrifugation steps. **Figure 2A** shows that we can remove methanol from the formulation by this process without losing too much Remdesivir. After just a single centrifuge, the resuspended nanocrystals were less than 0.002% methanol by volume and 4.9 mg/mL Remdesivir (initially 5.4 mg/mL). The size is about 450 nm after the centrifugation and resuspension (**Figure 2B**). Once methanol was removed, the formulations were frozen in liquid nitrogen with 20% trehalose (w/v) as a cryoprotectant where they could be stored long term. After thawing there was a modest increase in the nanocrystal size when thawed between 1 and 42 days after synthesis (**Figure 2C**). Suspension size was measured at 0 hrs and 4 hrs after thawing and there was a trend of increasing size over this time, but no changes in PDI were observed. In comparison to lyophilization (**Figure 2B**), the initial sizes were smaller when frozen rather than lyophilized. However, no further growth occurred during the 4 hours post-hydration when lyophilized. We concluded that both freezing and lyophilization were valid methods for long term storage of these suspensions, though they achieved slightly different terminal nanocrystal sizes. We opted to proceed using the freeze/thaw technique for long-term storage. Size increases were modest over 4 hours, and the crystals were of smaller size when frozen and thawed 550-600 nm compared to 750-950 nm when lyophilized. To thaw, nanocrystals were left at room temperature then shaken gently every few minutes for 15 min. At this point, the nanocrystal size was stable and nanocrystal suspension was ready for size measurements and to use for aerosol and other studies. To bypass the centrifugation and lyophilization steps, we also assessed the use of ethanol as an organic solvent for nanocrystal preparation by injecting Remdesivir into the aqueous media. Ethanol is already approved in aerosol formulations so complete removal is not as critical. Unfortunately, ethanol caused the nanocrystals to precipitate quickly and irreversibly, perhaps due to increased hydrophobic interactions, so ethanol was not explored further as a solvent for the nanocrystal preparation.

**Figure 2:**
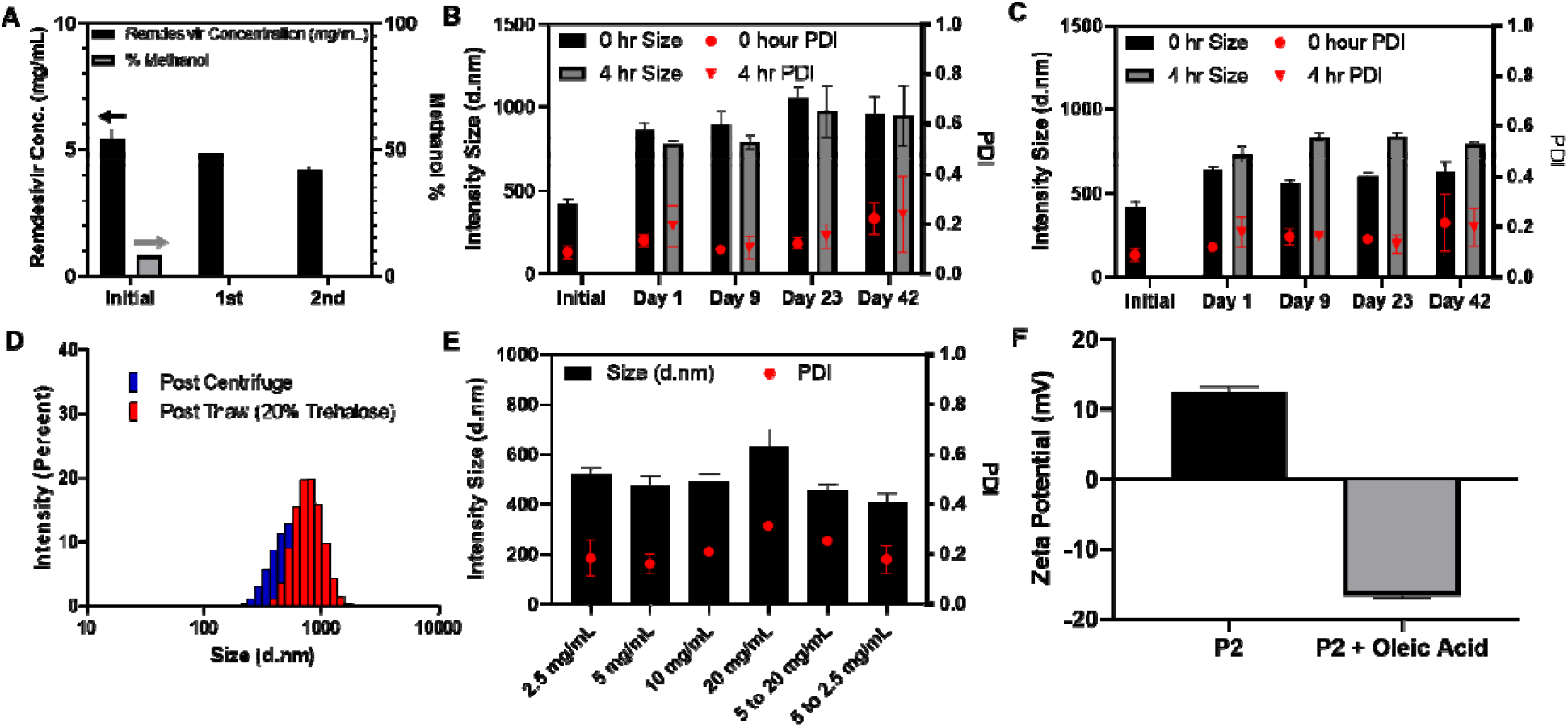
Reproducible preparation of stable, methanol-free P2 stabilized Remdesivir nanocrystal suspensions: **(A)** The concentration of Remdesivir (mg/ml) and the amount of methanol (volume %) remaining in the suspension after zero, one, and two centrifugal separations and redispersion of the suspension as measured by HPLC (Remdesivir) and ^1^H-NMR (methanol). Initial methanol volume % was 8.0% and after a single centrifuge was less than 0.002%. **(B)** The size (diameter) and PDI of P2 nanocrystals (0.25 mg/mL P2, 5 mg/mL Remdesivir after lyophilization in 20% trehalose (w/v) and stored at ambient temperature and hydrated with ultrapure water over 1-42 days after lyophilization. Sizes shown at 0 hr and 4 hr after hydration. **(C)** The size (diameter) and PDI of P2 nanocrystals (0.25 mg/mL P2, 5%) after a single centrifugation and freeze/thaw cycle in 20% Trehalose (v/v) over 1-42 days after freezing. Sizes shown at 0 hr and 4 hr after thaw. **(D)** The intensity size distribution of P2 nanocrystals (0.25 mg/mL P2, 5 mg/mL Remdesivir) after centrifugal removal of methanol and again after a freeze/thaw cycle in 20% trehalose (w/v). **(E)** The size of P2 nanocrystals after freeze/thaw (20% trehalose (w/v)) (0.125-1 mg/mL P2) when prepared at various Remdesivir concentrations (2.5-20 mg/mL Remdesivir) or when concentrated after centrifugation (5 mg/mL to 20 mg/mL) and then frozen or diluted (5 mg/mL to 2.5 mg/mL) and then frozen **(F)** The zeta potential of P2 (0.25 mg/mL) nanocrystals (5 mg/mL Remdesivir) with and without the addition of 0.005 mg/mL oleic acid as additional stabilizer.

Additionally, **Figure 2E** shows how the concentration of Remdesivir at synthesis does not affect the terminal size of the nanocrystals after freeze/thaw. We prepared crystals at concentrations ranging from 2.5 mg/mL to 20 mg/mL and the sizes remained in a narrow range from 475 to 632 nm. The PDI was slightly increased at the 20 mg/mL concentration, so we tested out ability to concentrate and dilute after a 5 mg/mL synthesis. Sizes and PDI were reduced when concentrating or diluting after centrifugation and prior to freezing. For certain applications, the addition of charged stabilizers to confer zeta potential to the particles could be advantageous. To this end, we added a small amount (0.005 mg/mL, 5 mg/mL Remdesivir) oleic acid as an additional stabilizer. Oleic Acid was chosen as it is an excipient in a clinically approved formulation for pulmonary delivery^36^. While it did not change the terminal size after freeze/thaw (**Supplemental Figure S4**), it did confer a significant negative zeta potential to the nanocrystals (**Figure 2F**). P2 stabilized nanocrystals had a positive zeta potential, while the addition of oleic acid rendered them negatively charged. Altering the particle surface charge and zeta potential in the future could be potentially useful for improved stability, changing nanoparticle binding with/penetration of mucus, and ultimately internalization into cells like type II pneumocytes which play a critical role in COVID-19 pathogenesis^50–52^. This introduces a degree of tunability to the particles and room for optimization *in vivo* as needed.

Once we established a final formulation of P2 stabilized nanocrystals at the 5% stabilizer relative to Remdesivir ratio (at any Remdesivir concentration) using stir bar mixing at room temperature, we wanted to evaluate some of their *in vitro* characteristics. Most nebulizers can hold up to 6 mL of formulation. At 10-20 mg doses, we would need roughly 1.5 to 5 mg/mL concentrations to deliver adequate drug, by our estimates of the requisite dosing (5-10% of IV dose). For our *in vitro* drug release study, we elected to use PBS at 37°C under sink conditions. Albumin, a major drug binding component of plasma, is not a major component of pleural fluid in the lungs making it unnecessary to include it in the release medium^53^. To compare, we also prepared a SBECD based cyclodextrin IC to mimic Gilead’s commercial Remdesivir formulation^8^. Gilead’s formulation contains a 1:30 ratio of Remdesivir to SBECD stabilizer at 5 mg/mL Remdesivir concentration (150 mg/mL SBECD, 96.8% excipient). Both formulations, nanocrystal and IC, were analyzed for drug release at 2 and 5 mg/mL initial concentrations (**Figure 3A**). All formulations had nearly identical drug release kinetic profiles indicating that the nanocrystals, like the ICs, are simple solubilizing formulations. However, the nanocrystals can solubilize Remdesivir at much lower excipient concentration than the SBECD based formulation (5% formulation weight POx compared to 96.7% formulation weight SBECD).

**Figure 3:**
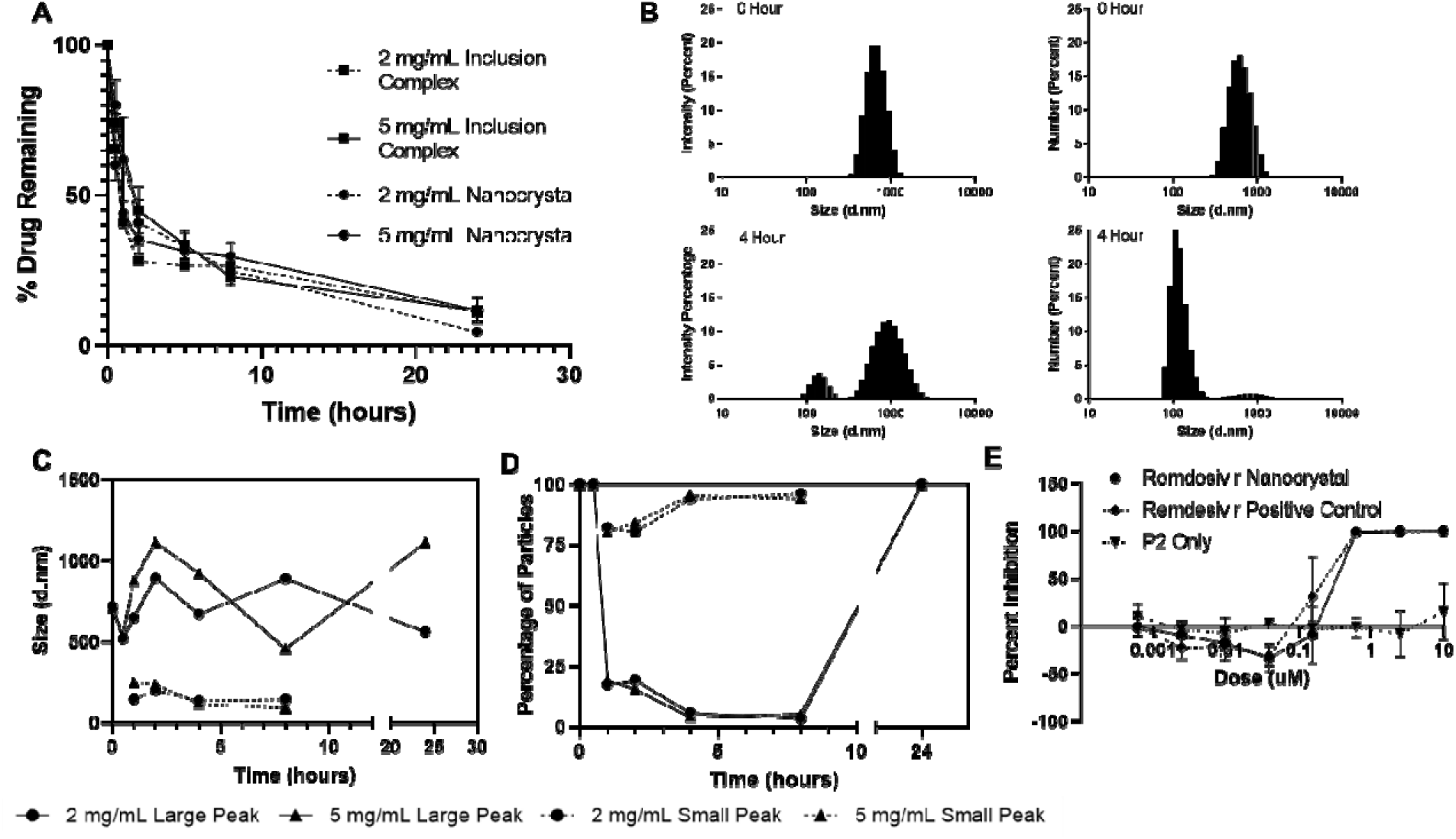
*In vitro* analysis of drug release from P2 stabilized Remdesivir nanocrystals:**(A)** The release of the drug over time in PBS at 37°C of nanocrystals at two concentrations and Remdesivir cyclodextrin inclusion complexes at the same concentrations. **(B)** The intensity and number distribution by DLS of nanocrystals at zero time point and after 4 hours of drug release. **(C)** The particle size corresponding to the two peaks observed in DLS intensity distribution plots at different time points during drug release process. **(D)** The percentage of particles that are in each peak in plot **(C)** as measured by the number distribution on DLS. **(E)** *In vitro* antiviral activity of the Remdesivir nanocrystals compared to Remdesivir positive control in DMSO and the P2 polymer stabilizer alone in the A549-Ace2 cell model^28^ **(A, C, D)** The drug concentration in the release experiment and size measurement was either 2 mg/ml or 5 mg/ml (0.10 or 0.25 mg/mL P2, respectively).

We also measured the changes in the particle size in the process of the drug release from the P2 stabilized nanocrystals. The initial nanocrystals at both drug concentrations had approximately the same size, about 750 nm (**Figure 3B** (top)). As time went on, the nanocrystals began to separate into smaller nanoparticles of roughly 150-200 nm as seen in **Figure 3B**, with the initial nanocrystals disappearing. Notably, on the number size distribution in **Figure 3B**, almost the entire particle population consists of the smaller 150-200 nm particles at 4 hours. This smaller population first appeared after around 1 hour of drug release and encompassed about 80% of the crystals present (**Figure 3C** and **3D**). Up until 8 hours, this population grew until it encompassed 95% of all crystals in solution. After 24 hours, when just about 10% of the drug remained in both the nanocrystal and SBECD formulations, the population had reverted to just a single peak. We hypothesize that upon dilution, the 750 nm nanocrystals break up into smaller aggregates which have a larger surface area/volume ratio. At later time points, only the larger crystals remain as most of the smaller ones have dissolved.

After demonstrating that our nanocrystals have a similar release profile to a commercial Remdesivir IC formulation mimic, we checked to ensure that our nanocrystal formulation maintains anti-viral efficacy (IC50 Comparison). Virus replication was monitored using SARS-CoV-2 virus that expresses nanoluciferase in place of ORF7a. Nanoluciferase expression as measured by luminescence correlates with virus replication. Replacement of ORF7a with a reporter protein does not impact virus replication and allows for high throughput screening of compounds for antiviral activity (24, 32, 33). A549 cells (human alveolar epithelial cells) expressing Ace2, a prominent receptor for SARS-CoV-2 (PMID: 32142651), are pre-treated with formulations at varied concentrations followed by infection with the SARS-CoV-2 reporter virus. After 48 hours, nanoluciferase expression is assayed and normalized to expression observed in DMSO treated controls. **Figure 3E** demonstrates that our nanocrystals maintain similar efficacy to the control free Remdesivir making them suitable candidates for aerosol delivery directly to the lungs.

Finally, we proceeded to analyze the formulations’ ability to be aerosolized into respirable particles using an air jet nebulizer. This was chosen as opposed to a vibrating mesh nebulizer to better facilitate the delivery of a drug suspension. The one we selected, the Nebutech 8960, was selected as a simple, inexpensive device which is readily available to respiratory therapists in hospitals. It reflects the urgent need to adopt a simple and inexpensive solution for a potentially large number of patients. Using our 10 mg/mL frozen formulation, it was thawed and diluted to a concentration of 1.5 mg/mL Remdesivir. We analyzed the aerosol particle output of the nebulizer from 0 to 2 min. **Figure 4A** shows how the choice of excipient can have significant impact on the foaming of the solution in the nebulizer. Generally, foaming leads to inconsistent output over time and should be avoided. Our P2 stabilized nanocrystals had minimal foaming when compared to just water. On the other hand, significant foaming was evident (**Figure 4A (right)**) in the F127 stabilized nanocrystals led to the inconsistent output over time as shown in **Supplemental Figure S2**. Another factor effecting aerosol particle output is the solution viscosity. **Supplementary Figure S5** shows there was no major difference in viscosity comparing water to our 2.5 mg/mL Remdesivir nanocrystal suspension (0.125 mg/mL P2). After being placed in the nebulizer and aerosolized for two minutes the P2 stabilized nanocrystals exhibited little if any change in the particle size compared to the initial suspension or that incubated at room temperature. **Figure 4C** shows the distribution of drug mass in particles of various aerosolized particle diameter cutoff ranges. We aerosolized P2 stabilized nanocrystals prepared without and with oleic acid. Both formulations had MMAD near the respirable particle size cutoff near 5.3 um.

**Figure 4:**
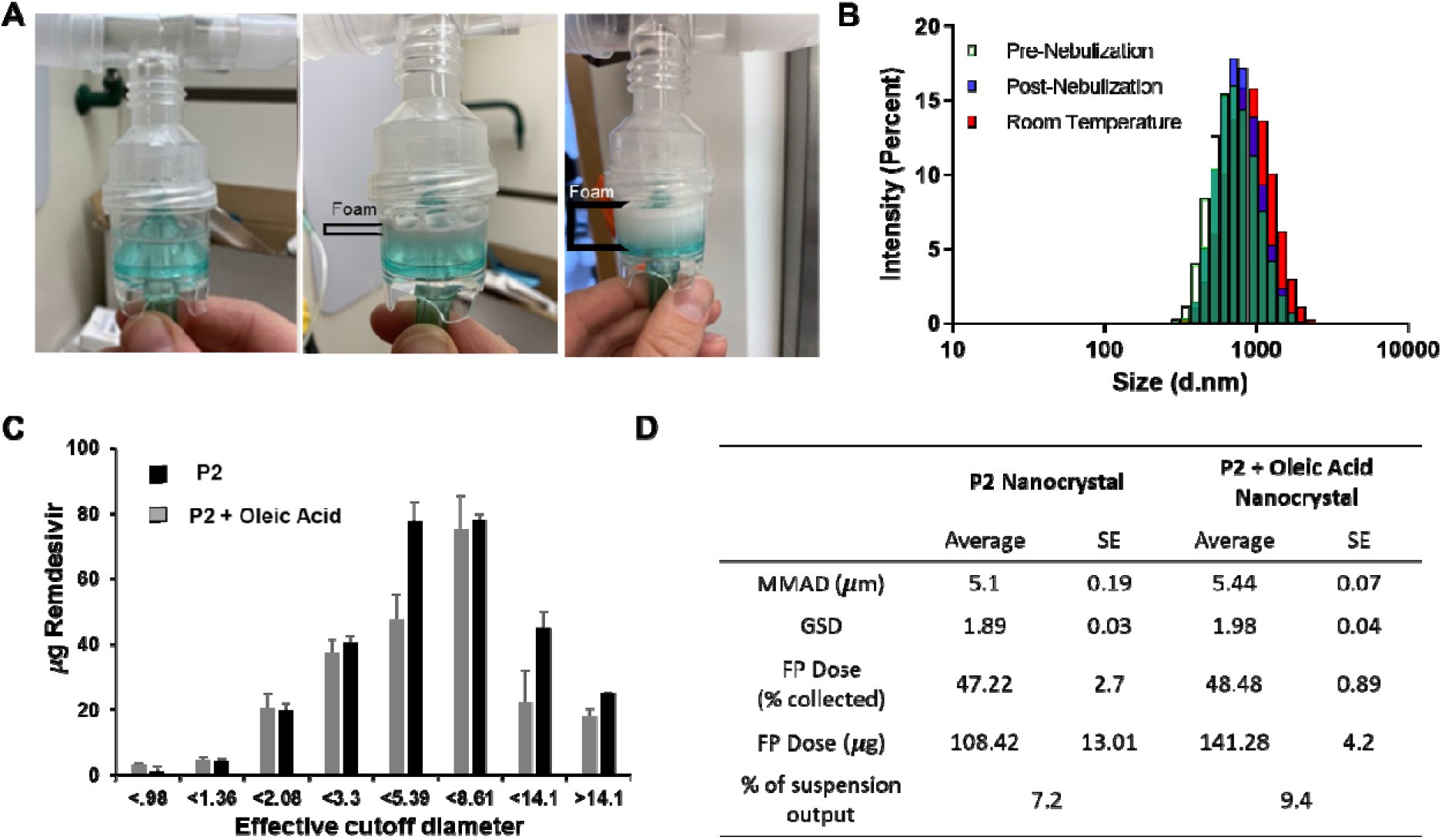
Aerosol output from nebulization of P2 stabilized nanocrystals. **(A)** The foaming of: left - water only; center - P2 stabilized nanocrystals; and (right) F127 stabilized nanocrystals (0.25 mg/mL stabilizer). **(B)** DLS particle size of P2 stabilized nanocrystals before (green) and after nebulization (blue), as well as after incubation at room temperature during the nebulization process (red). **(C)** Impactor analysis showing amount of Remdesivir mass in various particle cutoff sizes for P2 stabilized nanocrystals with and without 0.0015 mg/mL oleic acid (initial Remdesivir concentration at 1.5 mg/mL). Vacuum flow rate on NGI at 15 L/min, nanocrystal suspension volume at 3 mL, 2-minute collection time. (D) MMAD, GSD, % of Mass in fine particles (FP Dose %), FP Dose (ug), and percentage of nominal Remdesivir dose which was output are calculated as described in the methods.

In both formulations, the FP Dose % (fraction of mass that is in respirable particles) is near 50 %. The FP Dose in ug was 108.42 and 141.28 ug for the formulations prepared without and with oleic acid, respectively. The output rate is typically about 0.5 mL/minute for these nebulizers indicating that we delivered about 7.2 % and 9.4 % of the dose in the fine particles (respirable) using formulations prepared without and with oleic acid, respectively. These percentages compare the delivered dose in the fine particle fraction when compared to the nominal 1.5 mg/mL Remdesivir solution delivered over 2 minutes. Using the Impactor is not the most accurate measurement of total drug output, since there may be losses in sample recovery at each stage, so these measurements of drug output should be considered approximations which are likely lower than the true value.

## Discussion

It is clear that Remdesivir can prevent viral replication *in vitro*, but the success of the drug in humans has been called into question^10,11^. We believe this can be overcome, and Remdesivir use can be improved by altering the route of drug delivery. To this end, in late March 2020, we realized (along with others^54,55^) that an inhalable formulation of Remdesivir would be beneficial for COVID-19 treatment as it attacks the virus at the source of infection and avoids first-pass metabolism. COVID-19 disease progression begins in the upper respiratory tract, and it slowly progresses deeper into the lungs.

The SARS-CoV2 virus targets ciliated cells in the respiratory tract. Early data suggested that the infection begins in the nose, and then spreads deeper into the lungs, when the infection gets more serious^50^. Thus, delivering drug to the most serious site of infection, by inhalation delivery, would be ideal. Nebulizers (specifically jet nebulizers) are commonly used in clinical setting and are cheap, disposable, and easy to use. This solution-based drug product approach has a well understood development strategy and approval process with the FDA from a device perspective^33,34^. The absence of vaccines, and the high mortality rate, at the time this work was initiated made the prospects of rapid regulatory approval a priority in consideration of an appropriate delivery technology.

The formulation itself would require quick approval to be beneficial while vaccines remained elusive and as-yet untested with respect to long-term efficacy and necessity of a booster shot. Rapid approval of the formulation remains a priority and would be beneficial as new variants cause increasing concerns with disease transmissibility, severity, and vaccine efficacy. Criteria were developed to minimize barriers to the drug product formulation approval. These were: 1) achieve high solubility of Remdesivir (reduced overall inhaled dose volume from a nebulizer), 2) use GRAS or GRAS similar excipients, and 3) minimize the excipient use (ideally less than 5% of formulation weight) to maximize the drug for efficacy without impacting safety. To meet these requirements, we believed that a nanocrystal formulation could be the best approach for developing an inhalable Remdesivir therapy conducive to rapid approval. The bottom-up approach to nanocrystal synthesis is the simplest and involves the least expensive instruments and development process. This is perhaps more conducive to the rapid scale up and clinical approval. However, the bottom-up approach has its limitations. Namely, it can be more difficult to control crystal growth and produce small, monodisperse nanocrystals^27,35^.

If successful, an inhalable formulation of Remdesivir could produce higher Remdesivir concentrations at the site of disease without increasing systemic toxicity, which could potentially improve therapeutic outcomes in COVID-19. Indeed, there are several recent examples of nebulized treatments for COVID-19, some of which have entered clinical trials ([NCT05116865],[NCT04578236],[NCT04381364]). For example, Gilead has attempted to bring an inhaled formulation of Remdesivir to market [NCT04480333]. Their inhalable formulation is identical to their formulation designed for IV administration, utilizing SBECD at a 30:1 weight ratio to solubilize Remdesivir. The MMAD of the formulation was 2.7 um, and in a non-human primate model, they utilized a delivered dose (deposited in the lungs) of 0.35 mg/kg (over 60 minutes) to achieve lung concentrations of Remdesivir comparable to the 10 mg/K IV dose^24^. Thus, our initial assumption that dosing at 5-10% of the IV dose would be adequate was correct. Additionally, they utilized a Pari LC Plus nebulizer, which is of comparable design to our Nebutech 8960. Our approach delivers 7-9% of the formulation suspension concentration in “respirable particles” (<5 μm). In 10 kg non-human primates, we would require a dose of roughly 3.5 mg. We estimate that this dose could be delivered from our nanocrystal suspension over a comparable amount of time, 35 minutes (estimating 2.5 mg/mL Remdesivir suspension concentration, 8% respirable dose, 0.5 mL/min output rate). Utilizing a single excipient, at an extremely low concentration (Gilead: 150 mg/mL SBECD, P2 Nanocrystals: 0.125 mg/mL P2), we estimate we could deliver comparable doses to the SBECD-Remdesivir formulation in a shorter or comparable amount of time. In general, it is better to minimize excipient use, especially in sensitive organs like the lungs to minimize agitation and inflammation.

Other groups have also developed nanoparticle based formulations of Remdesivir as well. Vartak *et al.* have developed a liposomal formulation for inhalable delivery demonstrating an MMAD of 4.56 μm and an FPF of about 74% (P2 Nanocrystals ~50% FPF)^56^. However, this formulation still uses a significant amount of excipient and may be limited by drug release kinetics from the liposome or liposomal uptake into cells. Other liposomal formulations for inhalation have been developed with similar results^57^. Other systemic, PLGA formulations of Remdesivir have been developed as well^58^.

Notably, there is one clinically approved nanocrystal suspension on the market. It is a steroid formulation of the drug Budesonide for treating asthma developed by Astrazeneca (marketed as Pulmicort Respules ®)^50^. An *in vitro* analysis was performed on this formulation testing the influence of nebulizer choice, which greatly affects the delivered dose^60^. Additionally, FDA disclosure documents indicate that in *in vitro* conditions, only 17% of the nominal dose is actually delivered at the mouth piece-so this is our baseline delivery efficiency to achieve with our Remdesivir formulation. It is not clear if this 17% is in the fine particle fraction or a total output. In our formulation, we estimate that we deliver 15 % of the nominal concentration in the P2 Nanocrystals and 20 % in the P2-Oleic Acid Nanocrystals (based off total output, not just fine particle). Our formulation is close to reaching the standard set by the lone clinically approved nanocrystal suspension for aerosol delivery. Notably, both our formulation and the Budesonide suspension are designed to be used with jet-nebulizers. The nominal dose of Remdesivir which we need to deliver is significantly higher than Budesonide (Budesonide Dose=250-500 ug). To achieve higher doses, we can increase the concentration of the drug nanocrystals or at lower concentrations, nebulizers could easily be refilled and additional dose delivered.

There are many polymers which have been used to stabilize nanocrystals in the past such as Poloxamers (also known as “Pluronics”), which contain PEG, a GRAS agent. Additionally, some excipients are already clinically approved to be included in pulmonary formulations, such as ethanol, Tyloxapol, and oleic acid^36^. To the best of our knowledge, we are the first to report on the use of POx block copolymers for drug nanocrystal formulations, and in particular, inhalable drug delivery systems. As polymeric micelles, POx polymers have consistently demonstrated the ability to widen the therapeutic windows and improve therapeutic efficacy^25,47,61^. Notably, our P2 POx polymer was able to stabilize the drug nanocrystal of Remdesivir as the sole stabilizing agent and at very low concentrations. This was the only excipient we studied capable of this kind of stabilization under our stringent requirements. As POx polymers have shown the ability to solubilize a diverse array of poorly soluble drugs as micelles^25,48,61,62^, we believe they can also stabilize a vast array as nanocrystal formulations as well. While this is a step forward in improving therapy for COVID-19, it is perhaps more important in its implications as a semi-ubiquitous solubilizer for nanocrystal formulations which could be used to rapidly develop new drug formulations for existing and novel diseases.

## Conclusions

To conclude, we have developed a novel formulation of Remdesivir as a drug nanocrystal which is 95% Remdesivir by weight and conducive to the delivery of the drug directly to the site of disease, in the lungs. Remdesivir is highly efficacious against COVID-19 *in vitro*, but some questions have been raised about its *in vivo* efficacy. Roughly 50% of our formulation is nebulized into respirable particles (<5 um) and the output is consistent with the clinically approved Pulmicort Respules ® formulation from Astrazeneca-the only clinically approved drug nanocrystal suspension for pulmonary delivery. Our future work will focus on developing novel POx based excipients which could reduce drug nanocrystal size and lower the MMAD to improve delivery efficiency deep into the lungs. We believe that our POx platform could be used to rapidly develop nanocrystal formulations for future diseases to improve therapeutic outcomes.

## Acknowledgements

TEM was performed by A. Shankar Kumbhar at the Chapel Hill Analytical and Nanofabrication Laboratory (CHANL), a member of the North Carolina Research Triangle Nanotechnology Network (RTNN), which is supported by the NSF (grant ECCS-1542015) as part of the National Nanotechnology Coordinated Infrastructure (NNCI). We would like to acknowledge Dr. Raplh Baric for his support and insights.

## Data Availability

The raw/processed data required to reproduce these findings cannot be shared at this time due to legal reasons.

## References

(1) Lai, C.-C.; Shih, T.-P.; Ko, W.-C.; Tang, H.-J.; Hsueh, P.-R. Severe Acute Respiratory Syndrome Coronavirus 2 (SARS-CoV-2) and Coronavirus Disease-2019 (COVID-19): The Epidemic and the Challenges. Int J Antimicrob Agents 2020, 55 (3), 105924. https://doi.org/10.1016/j.ijantimicag.2020.105924.

(2) Guo, Y.-R.; Cao, Q.-D.; Hong, Z.-S.; Tan, Y.-Y.; Chen, S.-D.; Jin, H.-J.; Tan, K.-S.; Wang, D.-Y.; Yan, Y. The Origin, Transmission. and Clinical Therapies on Coronavirus Disease 2019 (COVID-19) Outbreak – an Update on the Status. Military Medical Research 2020, 7 (1), 11. https://doi.org/10.1186/s40779-020-00240-0.

(3) Ñamendys-Silva, S. A. Respiratory Support for Patients with COVID-19 Infection. The Lancet Respiratory Medicine 2020, 8 (4), e18. https://doi.org/10.1016/S2213-2600(20)30110-7.

(4) Shi, H.; Han, X.; Jiang, N.; Cao, Y.; Alwalid, O.; Gu, J.; Fan, Y.; Zheng, C. Radiological Findings from 81 Patients with COVID-19 Pneumonia in Wuhan, China: A Descriptive Study. The Lancet Infectious Diseases 2020, 20 (4), 425–434. https://doi.org/10.1016/S1473-3099(20)30086-4.

(5) Guy, R. K.; DiPaola, R. S.; Romanelli, F.; Dutch, R. E. Rapid Repurposing of Drugs for COVID-19. Science 2020, 368 (6493), 829–830. https://doi.org/10.1126/science.abb9332.

(6) Boras, B.; Jones, R. M.; Anson, B. J.; Arenson, D.; Aschenbrenner, L.; Bakowski, M. A.; Beutler, N.; Binder, J.; Chen, E.; Eng, H.; Hammond, H.; Hammond, J.; Haupt, R. E.; Hoffman, R.; Kadar, E. P.; Kania, R.; Kimoto, E.; Kirkpatrick, M. G.; Lanyon, L.; Lendy, E. K.; Lillis, J. R.; Logue, J.; Luthra, S. A.; Ma, C.; Mason, S. W.; McGrath, M. E.; Noell, S.; Obach, R. S.; O’Brien, M. N.; O’Connor, R.; Ogilvie, K.; Owen, D.; Pettersson, M.; Reese, M. R.; Rogers, T. F.; Rosales, R.; Rossulek, M. I.; Sathish, J. G.; Shirai, N.; Steppan, C.; Ticehurst, M.; Updyke, L. W.; Weston, S.; Zhu, Y.; White, K. M.; García-Sastre, A.; Wang, J.; Chatterjee, A. K.; Mesecar, A. D.; Frieman, M. B.; Anderson, A. S.; Allerton, C. Preclinical Characterization of an Intravenous Coronavirus 3CL Protease Inhibitor for the Potential Treatment of COVID19. 12 2021. https://doi.org/10.1038/s41467-021-26239-2.

(7) Lapadula, G.; Bernasconi, D. P.; Bellani, G.; Soria, A.; Rona, R.; Bombino, M.; Avalli, L.; Rondelli, E.; Cortinovis, B.; Colombo, E.; Valsecchi, M. G.; Migliorino, G. M.; Bonfanti, P.; Foti, G.; Remdesivir-Ria Study Group. Remdesivir Use in Patients Requiring Mechanical Ventilation Due to COVID-19. Open Forum Infectious Diseases 2020, 7 (11), ofaa481. https://doi.org/10.1093/ofid/ofaa481.

(8) Beigel, J. H.; Tomashek, K. M.; Dodd, L. E.; Mehta, A. K.; Zingman, B. S.; Kalil, A. C.; Hohmann, E.; Chu, H. Y.; Luetkemeyer, A.; Kline, S.; Lopez de Castilla, D.; Finberg, R. W.; Dierberg, K.; Tapson, V.; Hsieh, L.; Patterson, T. F.; Paredes, R.; Sweeney, D. A.; Short, W. R.; Touloumi, G.; Lye, D. C.; Ohmagari, N.; Oh, M.-D.; Ruiz-Palacios, G. M.; Benfield, T.; Fätkenheuer, G.; Kortepeter, M. G.; Atmar, R. L.; Creech, C. B.; Lundgren, J.; Babiker, A. G.; Pett, S.; Neaton, J. D.; Burgess, T. H.; Bonnett, T.; Green, M.; Makowski, M.; Osinusi, A.; Nayak, S.; Lane, H. C.; ACTT-1 Study Group Members. Remdesivir for the Treatment of Covid-19 - Final Report. N Engl J Med 2020, 383 (19), 1813–1826. https://doi.org/10.1056/NEJMoa2007764.

(9) Rochwerg, B.; Agarwal, A.; Zeng, L.; Leo, Y.-S.; Appiah, J. A.; Agoritsas, T.; Bartoszko, J.; Brignardello-Petersen, R.; Ergan, B.; Ge, L.; Geduld, H.; Gershengorn, H. B.; Manai, H.; Huang, M.; Lamontagne, F.; Kanda, S.; Kawano-Dourado, L.; Kurian, L.; Kwizera, A.; Murthy, S.; Qadir, N.; Siemieniuk, R.; Silvestre, M. A.; Vandvik, P. O.; Ye, Z.; Zeraatkar, D.; Guyatt, G. Remdesivir for Severe Covid-19: A Clinical Practice Guideline. BMJ 2020, 370, m2924. https://doi.org/10.1136/bmj.m2924.

(10) Rubin, D.; Chan-Tack, K.; Farley, J.; Sherwat, A. FDA Approval of Remdesivir — A Step in the Right Direction. New England Journal of Medicine 2020, 383 (27), 2598–2600. https://doi.org/10.1056/NEJMp2032369.

(11) Trkulja, V. Remdesivir for COVID-19 Pneumonia: Still Undecided, but It Might All Be about Adequate Timing. Eur J Clin Pharmacol 2021, 77 (6), 935–937. https://doi.org/10.1007/s00228-020-03085-7.

(12) Early Remdesivir to Prevent Progression to Severe Covid-19 in Outpatients | NEJM https://www.nejm.org/doi/full/10.1056/NEJMoa2116846 (accessed 2022-01-06).

(13) Mahase, E. Covid-19: Molnupiravir Reduces Risk of Hospital Admission or Death by 50% in Patients at Risk, MSD Reports. BMJ 2021, 375, n2422. https://doi.org/10.1136/bmj.n2422.

(14) Khoo, S. H.; Fitzgerald, R.; Fletcher, T.; Ewings, S.; Jaki, T.; Lyon, R.; Downs, N.; Walker, L.; Tansley-Hancock, O.; Greenhalf, W.; Woods, C.; Reynolds, H.; Marwood, E.; Mozgunov, P.; Adams, E.; Bullock, K.; Holman, W.; Bula, M. D.; Gibney, J. L.; Saunders, G.; Corkhill, A.; Hale, C.; Thorne, K.; Chiong, J.; Condie, S.; Pertinez, H.; Painter, W.; Wrixon, E.; Johnson, L.; Yeats, S.; Mallard, K.; Radford, M.; Fines, K.; Shaw, V.; Owen, A.; Lalloo, D. G.; Jacobs, M.; Griffiths, G. Optimal Dose and Safety of Molnupiravir in Patients with Early SARS-CoV-2: A Phase I, Open-Label, Dose Escalating, Randomized Controlled Study. Journal of Antimicrobial Chemotherapy 2021, 76 (12), 3286–3295. https://doi.org/10.1093/jac/dkab318.

(15) Holman, W.; Holman, W.; McIntosh, S.; Painter, W.; Painter, G.; Bush, J.; Cohen, O. Accelerated First-in-Human Clinical Trial of EIDD-2801/MK-4482 (Molnupiravir), a Ribonucleoside Analog with Potent Antiviral Activity against SARS-CoV-2. Trials 2021, 22 (1), 561. https://doi.org/10.1186/s13063-021-05538-5.

(16) Painter, W. P.; Holman, W.; Bush, J. A.; Almazedi, F.; Malik, H.; Eraut, N. C. J. E.; Morin, M. J.; Szewczyk, L. J.; Painter, G. R. Human Safety, Tolerability, and Pharmacokinetics of Molnupiravir, a Novel Broad-Spectrum Oral Antiviral Agent with Activity against SARS-CoV-2. Antimicrobial Agents and Chemotherapy 65 (5), e02428–20. https://doi.org/10.1128/AAC.02428-20.

(17) Weinreich, D. M.; Sivapalasingam, S.; Norton, T.; Ali, S.; Gao, H.; Bhore, R.; Musser, B. J.; Soo, Y.; Rofail, D.; Im, J.; Perry, C.; Pan, C.; Hosain, R.; Mahmood, A.; Davis, J. D.; Turner, K. C.; Hooper, A. T.; Hamilton, J. D.; Baum, A.; Kyratsous, C. A.; Kim, Y.; Cook, A.; Kampman, W.; Kohli, A.; Sachdeva, Y.; Graber, X.; Kowal, B.; DiCioccio, T.; Stahl, N.; Lipsich, L.; Braunstein, N.; Herman, G.; Yancopoulos, G. D. REGN-COV2, a Neutralizing Antibody Cocktail, in Outpatients with Covid-19. New England Journal of Medicine 2021, 384 (3), 238–251. https://doi.org/10.1056/NEJMoa2035002.

(18) Weinreich, D. M.; Sivapalasingam, S.; Norton, T.; Ali, S.; Gao, H.; Bhore, R.; Xiao, J.; Hooper, A. T.; Hamilton, J. D.; Musser, B. J.; Rofail, D.; Hussein, M.; Im, J.; Atmodjo, D. Y.; Perry, C.; Pan, C.; Mahmood, A.; Hosain, R.; Davis, J. D.; Turner, K. C.; Baum, A.; Kyratsous, C. A.; Kim, Y.; Cook, A.; Kampman, W.; Roque-Guerrero, L.; Acloque, G.; Aazami, H.; Cannon, K.; Simón-Campos, J. A.; Bocchini, J. A.; Kowal, B.; DiCioccio, T.; Soo, Y.; Stahl, N.; Lipsich, L.; Braunstein, N.; Herman, G.; Yancopoulos, G. D.; Investigators, for the T. REGEN-COV Antibody Cocktail Clinical Outcomes Study in Covid-19 Outpatients; 2021; p 2021.05.19.21257469. https://doi.org/10.1101/2021.05.19.21257469.

(19) Baum, A.; Fulton, B. O.; Wloga, E.; Copin, R.; Pascal, K. E.; Russo, V.; Giordano, S.; Lanza, K.; Negron, N.; Ni, M.; Wei, Y.; Atwal, G. S.; Murphy, A. J.; Stahl, N.; Yancopoulos, G. D.; Kyratsous, C. A. Antibody Cocktail to SARS-CoV-2 Spike Protein Prevents Rapid Mutational Escape Seen with Individual Antibodies. Science 2020, 369 (6506), 1014–1018. https://doi.org/10.1126/science.abd0831.

(20) Yan, V. C.; Muller, F. L. Advantages of the Parent Nucleoside GS-441524 over Remdesivir for Covid-19 Treatment. ACS Med. Chem. Lett. 2020, 11 (7), 1361–1366. https://doi.org/10.1021/acsmedchemlett.0c00316.

(21) Goldman, J. D.; Lye, D. C. B.; Hui, D. S.; Marks, K. M.; Bruno, R.; Montejano, R.; Spinner, C. D.; Galli, M.; Ahn, M.-Y.; Nahass, R. G.; Chen, Y.-S.; SenGupta, D.; Hyland, R. H.; Osinusi, A. O.; Cao, H.; Blair, C.; Wei, X.; Gaggar, A.; Brainard, D. M.; Towner, W. J.; Muñoz, J.; Mullane, K. M.; Marty, F. M.; Tashima, K. T.; Diaz, G.; Subramanian, A. Remdesivir for 5 or 10 Days in Patients with Severe Covid-19. New England Journal of Medicine 2020, 383 (19), 1827–1837. https://doi.org/10.1056/NEJMoa2015301.

(22) Gabriele, M.; Puccini, P.; Lucchi, M.; Vizziello, A.; Gervasi, P. G.; Longo, V. Presence and Inter-Individual Variability of Carboxylesterases (CES1 and CES2) in Human Lung. Biochem Pharmacol 2018, 150, 64–71. https://doi.org/10.1016/j.bcp.2018.01.028.

(23) Oesch, F.; Fabian, E.; Landsiedel, R. Xenobiotica-Metabolizing Enzymes in the Lung of Experimental Animals, Man and in Human Lung Models. Arch Toxicol 2019, 93 (12), 3419–3489. https://doi.org/10.1007/s00204-019-02602-7.

(24) Vermillion, M. S.; Murakami, E.; Ma, B.; Pitts, J.; Tomkinson, A.; Rautiola, D.; Babusis, D.; Irshad, H.; Siegel, D.; Kim, C.; Zhao, X.; Niu, C.; Yang, J.; Gigliotti, A.; Kadrichu, N.; Bilello, J. P.; Ellis, S.; Bannister, R.; Subramanian, R.; Smith, B.; Mackman, R. L.; Lee, W. A.; Kuehl, P. J.; Hartke, J.; Cihlar, T.; Porter, D. P. Inhaled Remdesivir Reduces Viral Burden in a Nonhuman Primate Model of SARS-CoV-2 Infection. Science Translational Medicine 2021. https://doi.org/10.1126/scitranslmed.abl8282.

(25) Luxenhofer, R.; Han, Y.; Schulz, A.; Tong, J.; He, Z.; Kabanov, A. V.; Jordan, R. Poly(2-Oxazoline)s as Polymer Therapeutics. Macromolecular Rapid Communications 2012, 33 (19), 1613–1631. https://doi.org/10.1002/marc.201200354.

(26) Hwang, D.; Ramsey, J. D.; Makita, N.; Sachse, C.; Jordan, R.; Sokolsky-Papkov, M.; Kabanov, A. V. Novel Poly(2-Oxazoline) Block Copolymer with Aromatic Heterocyclic Side Chains as a Drug Delivery Platform. Journal of Controlled Release 2019, 307, 261–271. https://doi.org/10.1016/j.jconrel.2019.06.037.

(27) Sinha, B.; Müller, R. H.; Möschwitzer, J. P. Bottom-up Approaches for Preparing Drug Nanocrystals: Formulations and Factors Affecting Particle Size. International Journal of Pharmaceutics 2013, 453 (1), 126–141. https://doi.org/10.1016/j.ijpharm.2013.01.019.

(28) Hou, Y. J.; Chiba, S.; Halfmann, P.; Ehre, C.; Kuroda, M.; Dinnon, K. H.; Leist, S. R.; Schäfer, A.; Nakajima, N.; Takahashi, K.; Lee, R. E.; Mascenik, T. M.; Graham, R.; Edwards, C. E.; Tse, L. V.; Okuda, K.; Markmann, A. J.; Bartelt, L.; de Silva, A.; Margolis, D. M.; Boucher, R. C.; Randell, S. H.; Suzuki, T.; Gralinski, L. E.; Kawaoka, Y.; Baric, R. S. SARS-CoV-2 D614G Variant Exhibits Efficient Replication Ex Vivo and Transmission in Vivo. Science 2020, 370 (6523), 1464–1468. https://doi.org/10.1126/science.abe8499.

(29) Rappazzo, C. G.; Tse, L. V.; Kaku, C. I.; Wrapp, D.; Sakharkar, M.; Huang, D.; Deveau, L. M.; Yockachonis, T. J.; Herbert, A. S.; Battles, M. B.; O’Brien, C. M.; Brown, M. E.; Geoghegan, J. C.; Belk, J.; Peng, L.; Yang, L.; Hou, Y.; Scobey, T. D.; Burton, D. R.; Nemazee, D.; Dye, J. M.; Voss, J. E.; Gunn, B. M.; McLellan, J. S.; Baric, R. S.; Gralinski, L. E.; Walker, L. M. Broad and Potent Activity against SARS-like Viruses by an Engineered Human Monoclonal Antibody. Science 2021, 371 (6531), 823–829. https://doi.org/10.1126/science.abf4830.

(30) Hickey, A. Methods of Aerosol Particle Size Characterization. In Pharmaceutical Inhalation Aerosol Technology; Hickey, A., Ed.; Marcel Dekker, Inc., 2003; pp 345–384.

(31) Hickey, A. J. Emerging Trends in Inhaled Drug Delivery. Adv Drug Deliv Rev 2020, 157, 63–70. https://doi.org/10.1016/j.addr.2020.07.006.

(32) Hanafin, P. O.; Jermain, B.; Hickey, A. J.; Kabanov, A. V.; Kashuba, A. DM.; Sheahan, T. P.; Rao, G. G. A Mechanism-Based Pharmacokinetic Model of Remdesivir Leveraging Interspecies Scaling to Simulate COVID-19 Treatment in Humans. CPT: Pharmacometrics & Systems Pharmacology 2021, 10 (2), 89–99. https://doi.org/10.1002/psp4.12584.

(33) de Boer, A.; Thalberg, K. Devices and Fomulations: General Introduction and Wet Aerosol Delivery Systems. In Inhaled Medicines; Kassinos, S., Backman, P., Conway, J., Hickey, A., Eds.; Elsevier, 2021; pp 35–65. https://doi.org/10.1016/C2017-0-02102-X.

(34) Pritchard, J.; van Hollen, D.; Hatley, R. Nebulizers. In Pharmaceutical Inhalation Aerosol Technologies; Hickey, A., da Rocha, S., Eds.; CRC Press: Boca Raton, FL, 2019; pp 473–492.

(35) Van Eerdenbrugh, B.; Van den Mooter, G.; Augustijns, P. Top-down Production of Drug Nanocrystals: Nanosuspension Stabilization, Miniaturization and Transformation into Solid Products. International Journal of Pharmaceutics 2008, 364 (1), 64–75. https://doi.org/10.1016/j.ijpharm.2008.07.023.

(36) Pulmonary Drug Delivery: Advances and Challenges | Wiley https://www.wiley.com/en-us/Pulmonary+Drug+Delivery%3A+Advances+and+Challenges-p-9781118799543 (accessed 2021-12-14).

(37) Otsuka, H.; Nagasaki, Y.; Kataoka, K. PEGylated Nanoparticles for Biological and Pharmaceutical Applications. Advanced Drug Delivery Reviews 2003, 55 (3), 403–419. https://doi.org/10.1016/S0169-409X(02)00226-0.

(38) Mozar, F. S.; Chowdhury, E. H. Impact of PEGylated Nanoparticles on Tumor Targeted Drug Delivery. Current Pharmaceutical Design 2018, 24 (28), 3283–3296. https://doi.org/10.2174/1381612824666180730161721.

(39) Pitto-Barry, A.; E. Barry, N. P. Pluronic® Block-Copolymers in Medicine: From Chemical and Biological Versatility to Rationalisation and Clinical Advances. Polymer Chemistry 2014, 5 (10), 3291–3297. https://doi.org/10.1039/C4PY00039K.

(40) Ball, R. L.; Bajaj, P.; Whitehead, K. A. Achieving Long-Term Stability of Lipid Nanoparticles: Examining the Effect of PH, Temperature, and Lyophilization. Int J Nanomedicine 2016, 12, 305–315. https://doi.org/10.2147/IJN.S123062.

(41) Zhao, P.; Hou, X.; Yan, J.; Du, S.; Xue, Y.; Li, W.; Xiang, G.; Dong, Y. Long-Term Storage of Lipid-like Nanoparticles for MRNA Delivery. Bioactive Materials 2020, 5 (2), 358–363. https://doi.org/10.1016/j.bioactmat.2020.03.001.

(42) Almalik, A.; Alradwan, I.; Kalam, M. A.; Alshamsan, A. Effect of Cryoprotection on Particle Size Stability and Preservation of Chitosan Nanoparticles with and without Hyaluronate or Alginate Coating. Saudi Pharmaceutical Journal 2017, 25 (6), 861–867. https://doi.org/10.1016/j.jsps.2016.12.008.

(43) Chung, N.-O.; Lee, M. K.; Lee, J. Mechanism of Freeze-Drying Drug Nanosuspensions. International Journal of Pharmaceutics 2012, 437 (1), 42–50. https://doi.org/10.1016/j.ijpharm.2012.07.068.

(44) Louey, M. D.; Van Oort, M.; Hickey, A. J. Aerosol Dispersion of Respirable Particles in Narrow Size Distributions Using Drug-Alone and Lactose-Blend Formulations. Pharm Res 2004, 21 (7), 1207–1213. https://doi.org/10.1023/B:PHAM.0000032991.74736.3e.

(45) Kendrick, A. H.; Smith, E. C.; Wilson, R. S. Selecting and Using Nebuliser Equipment. Thorax 1997, 52 (Suppl 2), S92–S101.

(46) Hess, D.; Fisher, D.; Williams, P.; Pooler, S.; Kacmarek, R. M. Medication Nebulizer Performance: Effects Of Diluent Volume, Nebulizer Flow, and Nebulizer Brand. Chest 1996, 110 (2), 498–505. https://doi.org/10.1378/chest.110.2.498.

(47) Luxenhofer, R.; Schulz, A.; Roques, C.; Li, S.; Bronich, T. K.; Batrakova, E. V.; Jordan, R.; Kabanov, A. V. Doubly-Amphiphilic Poly(2-Oxazoline)s as High-Capacity Delivery Systems for Hydrophobic Drugs. Biomaterials 2010, 31 (18), 4972–4979. https://doi.org/10.1016/j.biomaterials.2010.02.057.

(48) Hwang, D.; Ramsey, J. D.; Kabanov, A. V. Polymeric Micelles for the Delivery of Poorly Soluble Drugs: From Nanoformulation to Clinical Approval. Advanced Drug Delivery Reviews 2020. https://doi.org/10.1016/j.addr.2020.09.009.

(49) Lim, C.; Ramsey, J. D.; Hwang, D.; Teixeira, S. C. M.; Poon, C.-D.; Strauss, J. D.; Sokolsky-Papkov, M.; Kabanov, A. V. Drug-Dependent Morphological Transitions in Spherical and Worm-like Polymeric Micelles Define Stability and Pharmacological Performance of Micellar Drugs; 2021; p 2021.06.10.447962. https://doi.org/10.1101/2021.06.10.447962.

(50) Hou, Y. J.; Okuda, K.; Edwards, C. E.; Martinez, D. R.; Asakura, T.; Dinnon, K. H.; Kato, T.; Lee, R. E.; Yount, B. L.; Mascenik, T. M.; Chen, G.; Olivier, K. N.; Ghio, A.; Tse, L. V.; Leist, S. R.; Gralinski, L. E.; Schäfer, A.; Dang, H.; Gilmore, R.; Nakano, S.; Sun, L.; Fulcher, M. L.; Livraghi-Butrico, A.; Nicely, N. I.; Cameron, M.; Cameron, C.; Kelvin, D. J.; de Silva, A.; Margolis, D. M.; Markmann, A.; Bartelt, L.; Zumwalt, R.; Martinez, F. J.; Salvatore, S. P.; Borczuk, A.; Tata, P. R.; Sontake, V.; Kimple, A.; Jaspers, I.; O’Neal, W. K.; Randell, S. H.; Boucher, R. C.; Baric, R. S. SARS-CoV-2 Reverse Genetics Reveals a Variable Infection Gradient in the Respiratory Tract. Cell 2020, 182 (2), 429–446.e14. https://doi.org/10.1016/j.cell.2020.05.042.

(51) Müller, R. H.; Jacobs, C.; Kayser, O. Nanosuspensions as Particulate Drug Formulations in Therapy: Rationale for Development and What We Can Expect for the Future. Advanced Drug Delivery Reviews 2001, 47 (1), 3–19. https://doi.org/10.1016/S0169-409X(00)00118-6.

(52) Schattling, P.; Taipaleenmäki, E.; Zhang, Y.; Städler, B. A Polymer Chemistry Point of View on Mucoadhesion and Mucopenetration. Macromolecular Bioscience 2017, 17 (9), 1700060. https://doi.org/10.1002/mabi.201700060.

(53) Veldhuizen, E. J. A.; Haagsman, H. P. Role of Pulmonary Surfactant Components in Surface Film Formation and Dynamics. Biochimica et Biophysica Acta (BBA) - Biomembranes 2000, 1467 (2), 255–270. https://doi.org/10.1016/S0005-2736(00)00256-X.

(54) Sahakijpijarn, S.; Moon, C.; Warnken, Z. N.; Maier, E. Y.; DeVore, J. E.; Christensen, D. J.; Koleng, J. J.; Williams, R. O. In Vivo Pharmacokinetic Study of Remdesivir Dry Powder for Inhalation in Hamsters. International Journal of Pharmaceutics: X 2021, 3, 100073. https://doi.org/10.1016/j.ijpx.2021.100073.

(55) Sahakijpijarn, S.; Moon, C.; Koleng, J. J.; Christensen, D. J.; Williams, R. O. Development of Remdesivir as a Dry Powder for Inhalation by Thin Film Freezing. Pharmaceutics 2020, 12 (11), 1002. https://doi.org/10.3390/pharmaceutics12111002.

(56) Vartak, R.; Patil, S. M.; Saraswat, A.; Patki, M.; Kunda, N. K.; Patel, K. Aerosolized Nanoliposomal Carrier of Remdesivir: An Effective Alternative for COVID-19 Treatment in Vitro. Nanomedicine 2021, 16 (14), 1187–1202. https://doi.org/10.2217/nnm-2020-0475.

(57) Li, J.; Zhang, K.; Wu, D.; Ren, L.; Chu, X.; Qin, C.; Han, X.; Hang, T.; Xu, Y.; Yang, L.; Yin, L. Liposomal Remdesivir Inhalation Solution for Targeted Lung Delivery as a Novel Therapeutic Approach for COVID-19. Asian Journal of Pharmaceutical Sciences 2021, 16 (6), 772–783. https://doi.org/10.1016/j.ajps.2021.09.002.

(58) Patki, M.; Palekar, S.; Reznik, S.; Patel, K. Self-Injectable Extended Release Formulation of Remdesivir (SelfExRem): A Potential Formulation Alternative for COVID-19 Treatment. Int J Pharm 2021, 597, 120329. https://doi.org/10.1016/j.ijpharm.2021.120329.

(59) Karlsson, A.-K.; SE; Larrivee-Elkins, C.; Molin, O.; SE. United States Patent: 7524834 - Sterile Powders, Formulations, and Methods for Producing the Same. 7524834, April 28, 2009.

(60) Barry, P. W.; O’Callaghan, C. An in Vitro Analysis of the Output of Budesonide from Different Nebulizers. Journal of Allergy and Clinical Immunology 1999, 104 (6), 1168–1173. https://doi.org/10.1016/S0091-6749(99)70009-6.

(61) Wan, X.; Beaudoin, J. J.; Vinod, N.; Min, Y.; Makita, N.; Bludau, H.; Jordan, R.; Wang, A.; Sokolsky, M.; Kabanov, A. V. Co-Delivery of Paclitaxel and Cisplatin in Poly(2-Oxazoline) Polymeric Micelles: Implications for Drug Loading, Release, Pharmacokinetics and Outcome of Ovarian and Breast Cancer Treatments. Biomaterials 2019, 192, 1–14. https://doi.org/10.1016/j.biomaterials.2018.10.032.

(62) Alves, V. M.; Hwang, D.; Muratov, E.; Sokolsky-Papkov, M.; Varlamova, E.; Vinod, N.; Lim, C.; Andrade, C. H.; Tropsha, A.; Kabanov, A. Cheminformatics-Driven Discovery of Polymeric Micelle Formulations for Poorly Soluble Drugs. Science Advances 2019, 5 (6), eaav9784. https://doi.org/10.1126/sciadv.aav9784.

