## Supplemental Data for "Nanoformulated Remdesivir with Extremely Low Content of Poly(2-oxazoline) - Based Stabilizer for Aerosol Treatment of COVID-19"

**Supplemental Table S1:** Excipients evaluated in this work
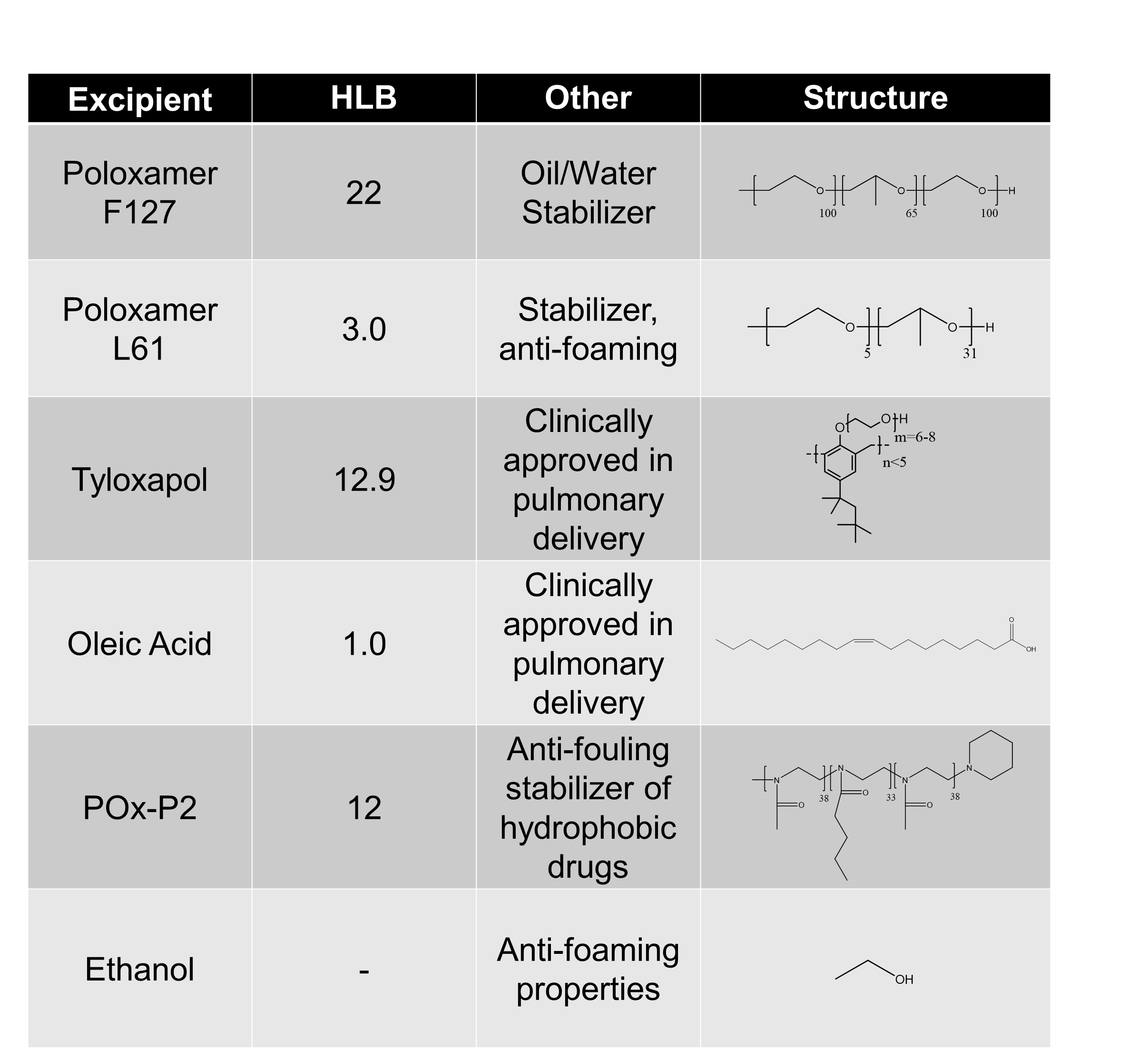

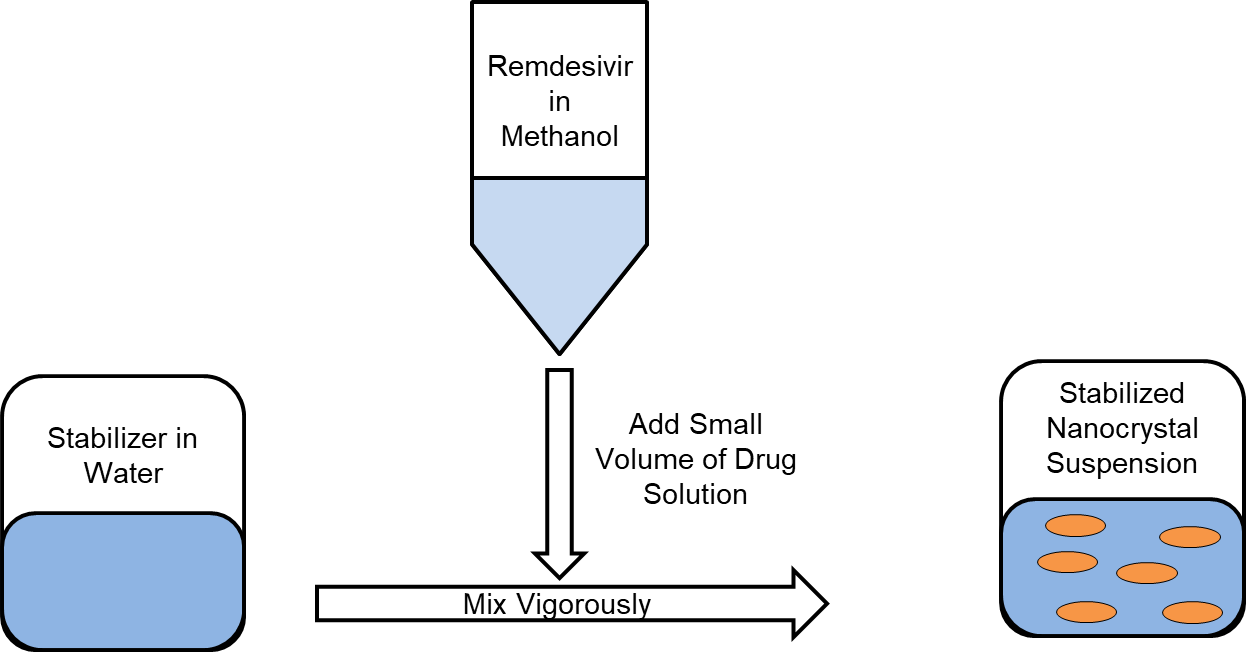

**Supplemental Figure S1:** Synthetic scheme of F127 stabilized Remdesivir nanocrystals

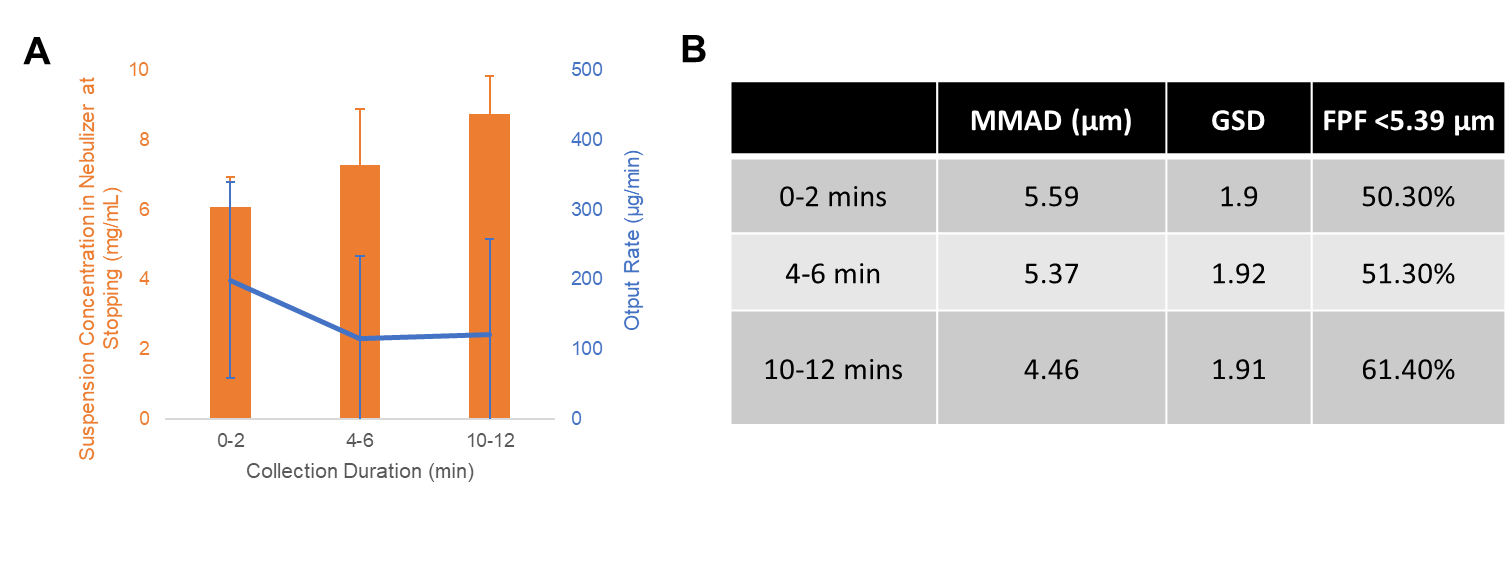

**Supplemental Figure S2:** Aerosol output of 0.25 mg/mL F127 Remdesivir (5 mg/mL) nanocrystals: **(a)** the nanocrystals suspension concentration and the output rate of drug over time and **(b)** The MMAD, GSD, and FPF of the particles over the nebulizer output time range.

**Supplemental Table S2:** Select excipient combinations and the nanocrystal size(DLS Intensity Distribution)/stability of the produced nanocrystals (5 mg/mL Remdesivir) when left at room temperature.

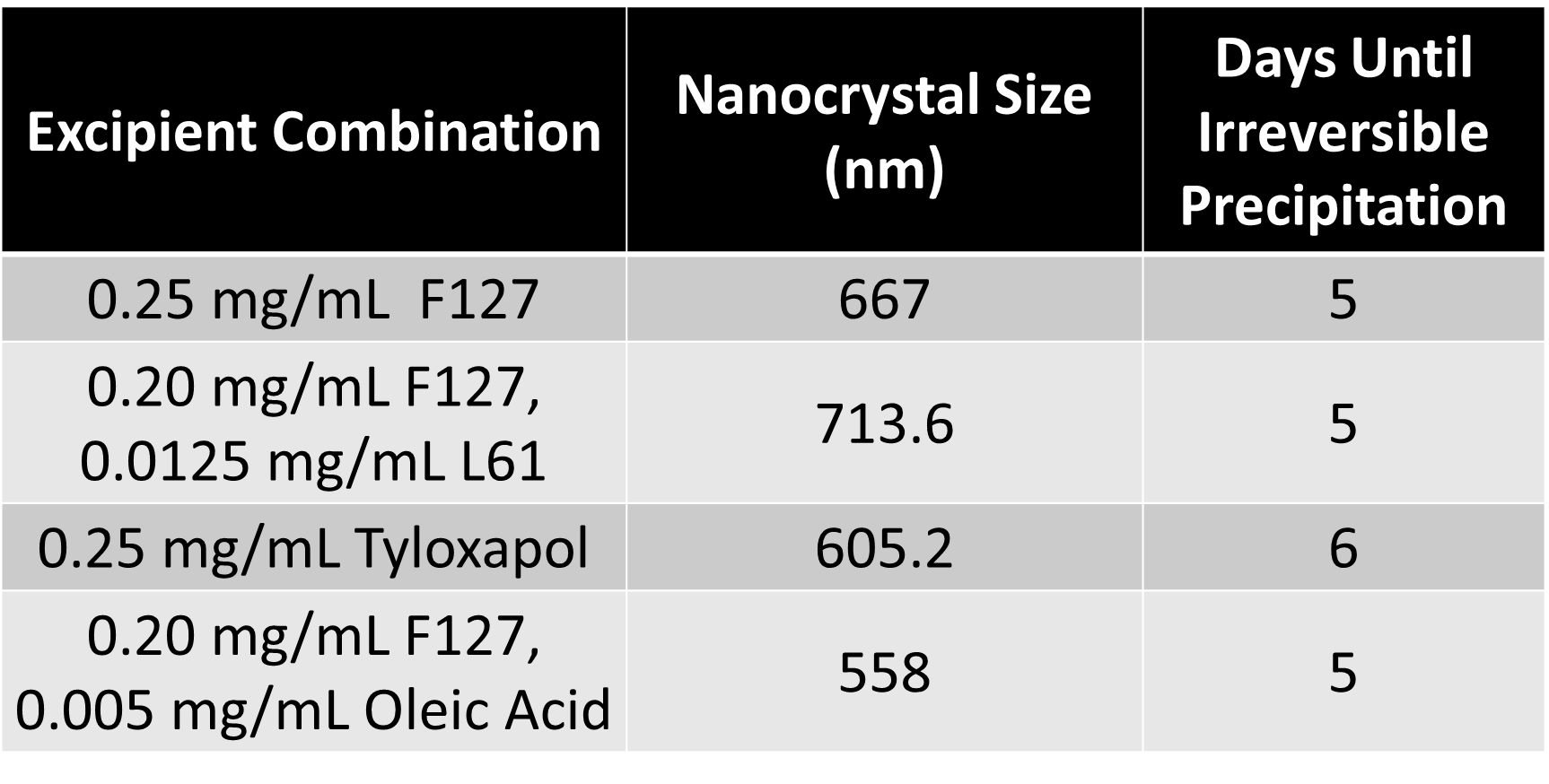

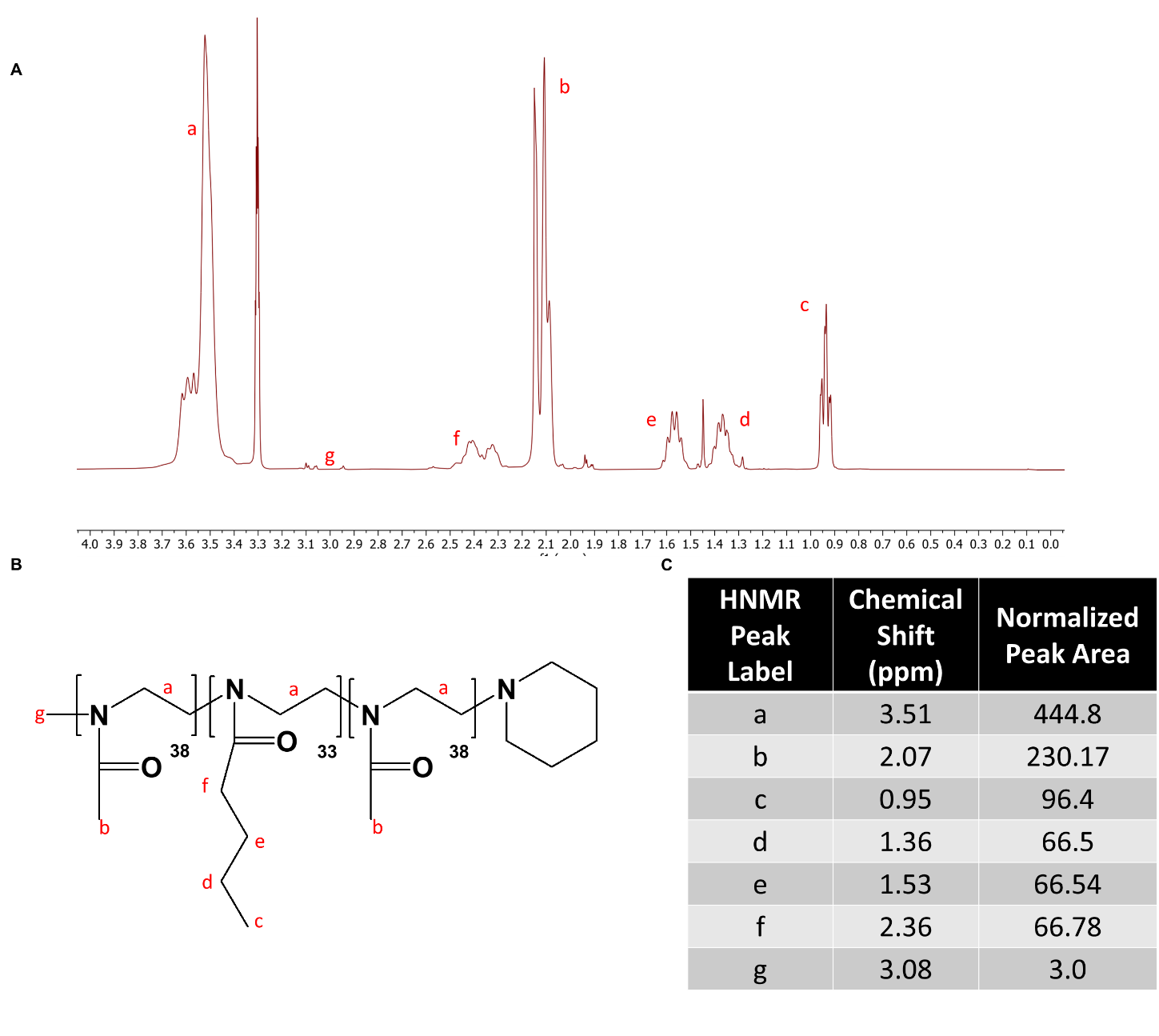

**Supplemental Figure S3: (A)** ^1^H-NMR spectrum and **(B)** Structure with **(C)** block length confirmation of P2 polymer.

**Supplemental Table S3:** Nanocrystal stability after multiple centrifugations at various speeds, DLS Size, RDV concentration. PC1 (Post-Centrifuge 1) is the pellet suspended in water after the first centrifugation. PC2 (Post-Centrifuge 2) is the pellet suspended in water after the second centrifugation. Starting nanocrystal concentration was 4 mg/mL Remdesivir, 0.2 mg/mL P2 stabilizer.

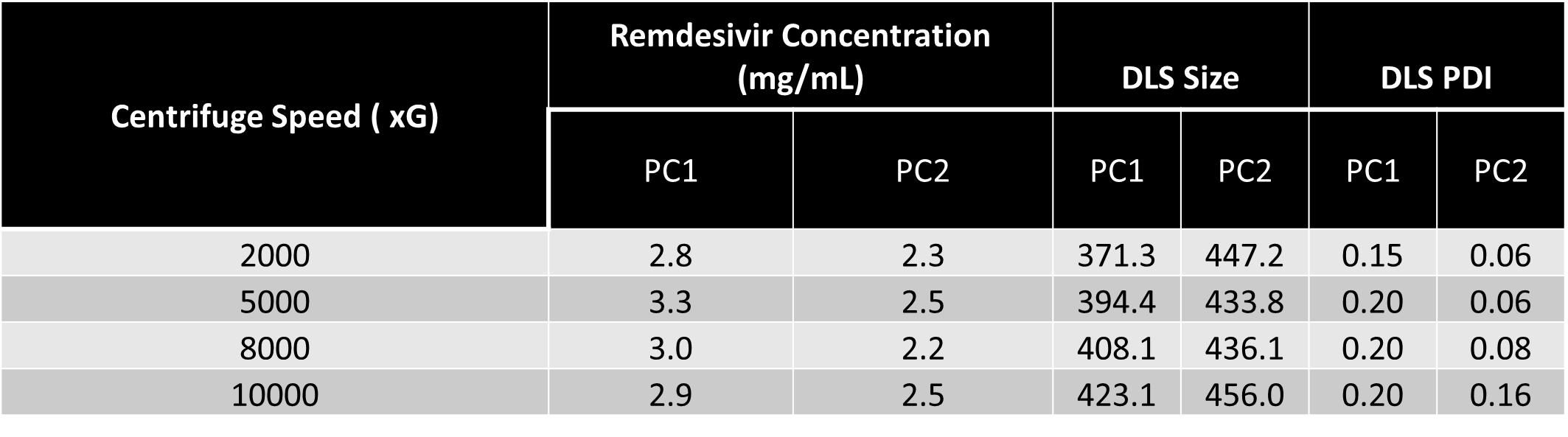

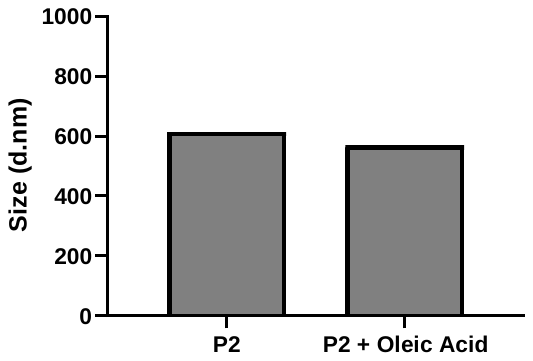

**Supplemental Figure S4:** Oleic Acid inclusion at 0.005 mg/mL does not change the size of Remdesivir Nanocrystals after a freeze/thaw cycle in 20% Trehalose (DLS intensity distribution)

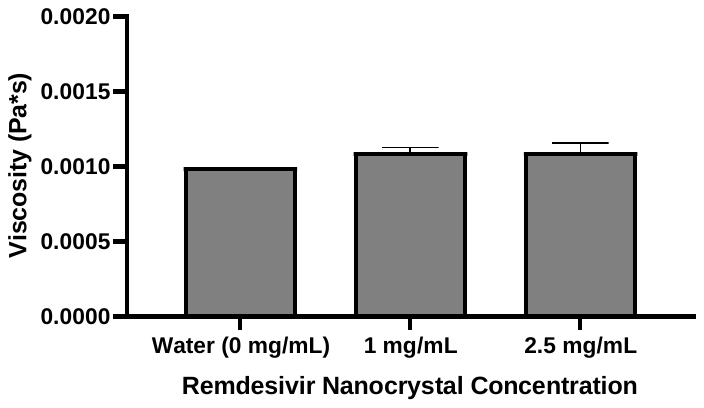

**Supplemental Figure S5:** Viscosity of Remdesivir NC formulations compared to pure water. Viscosity values are not statistically different.
